# A novel Mycobacterium tuberculosis-specific subunit vaccine provides synergistic immunity upon co-administration with Bacillus Calmette-Guérin

**DOI:** 10.1101/2021.03.18.435784

**Authors:** Joshua S. Woodworth, Helena Strand Clemmensen, Hannah Battey, Karin Dijkman, Thomas Lindenstrøm, Raquel Salvador Laureano, Randy Taplitz, Jeffrey Morgan, Claus Aagaard, Ida Rosenkrands, Cecilia S. Lindestam Arlehamn, Peter Andersen, Rasmus Mortensen

## Abstract

Tuberculosis (TB) remains a global health crisis. Following encouraging clinical results of subunit vaccination and revaccination with Bacillus Calmette-Guérin (BCG), it has been suggested to combine BCG and subunit vaccines for increased efficacy. Current subunit vaccines are almost exclusively designed as BCG boosters. The goal of this study was to design a subunit vaccine that does not share antigens with BCG and explore the advantages of a BCG+subunit vaccine co-administration strategy, where the two vaccines do not cross-react. Eight protective antigens were selected to create a Mycobacterium tuberculosis (Mtb)-specific subunit vaccine, named H107. Whereas subunit vaccines with BCG-shared antigens displayed cross-reactivity to BCG in vivo in both mice and humans, H107 showed no cross-reactivity and did not inhibit BCG colonization in mice. Encouragingly, co-administering H107 with BCG (BCG+H107) led to increased adaptive immune responses against both H107 and BCG leading to improved BCG-mediated immunity. In contrast to subunit vaccines with BCG-shared antigens, ‘boosting’ BCG with H107 led to substantial expansion of clonal diversity in the T cell repertoire, and BCG+H107 co-administration conferred significantly increased Th17 responses and less differentiated CD4 T cells. CD4 T cells induced by BCG+H107 maintained functional superiority after Mtb infection, and BCG+H107 provided significantly increased long-term protection compared to both BCG and H107 alone, as well as, BCG co-administered with a subunit vaccine composed of antigens shared with BCG. Overall, we identify several advantages of combining an Mtb-specific subunit vaccine with BCG and introduce H107 as a BCG-complementing vaccine with distinctive value for co-administration with BCG.

## INTRODUCTION

Despite decades of mass vaccination with the current tuberculosis (TB) vaccine, Bacillus Calmette-Guérin (BCG), *Mycobacterium tuberculosis* (Mtb), the primary causative agent of TB, remains one of the deadliest infectious agents (*1*). Due to the impact of the ongoing COVID-19 pandemic on TB case-finding and treatment, the number of cases is expected to rise significantly over the coming years (*2*). Taken together with the increasing incidence of multi-drug resistant TB (MDR-TB) strains (*3*), Mtb poses one of the greatest challenges for global health in the future, and only with a new and more efficacious TB vaccine strategy can the TB epidemic be ended.

Primary neonatal immunization with BCG does not reliably prevent pulmonary TB in adults and adolescents (*4*), and several studies investigating revaccination of adolescents and adults with BCG to boost immunity have shown little or no added protection (*5–8*). Geographical variations in environmental mycobacteria are associated with variable efficacy after BCG revaccination (*9–11*). As a live attenuated vaccine, BCG is dependent upon replication and/or persistence to confer immunity and protection, and immunological sensitization via exposure to non-tuberculous mycobacteria (NTM) has been shown to induce cross-reacting immune responses that can inhibit BCG replication and decrease efficacy (*9, 12*). However, a recent report showed a vaccine efficacy (VE) of BCG revaccination of 45.4% against sustained Mtb infection in a cohort of 660 previously uninfected South African adolescents (*13*). There is therefore renewed optimism that this approach can have success in some geographical regions, and a large confirmatory study of BCG revaccination with 1,800 uninfected adolescents across several countries in southern Africa is currently underway (NCT04152161).

In contrast to BCG and other live vaccines, non-replicating subunit vaccines rely on synthetic immunostimulatory adjuvants and are therefore refractory to prior mycobacterial sensitization. Adjuvanted protein subunit vaccines also hold the highest benefit of safety, which is of particular importance given the high occurrence of TB in HIV-prevalent settings (*1*). Encouragingly, two subunit vaccines have recently demonstrated the first signals of VE in clinical trials: H4:IC31® (VE 30.5%) (*13*) and M72/AS01E (VE 49.7%) (*14*). Despite these promising results, vaccines with improved efficacy of >50% are considered necessary to reach the target of the World Health Organization (WHO) End TB strategy (*15*) and there has been a call for new vaccine candidates that broaden the antigen repertoire towards this endeavor (*16–18*).

Based on the encouraging breakthroughs in TB vaccine clinical development, it has been suggested that subunit vaccines could be given in combination with BCG (re)vaccination to create a more efficacious vaccine regimen (*19*). Such a strategy would be in accordance with WHO’s recommendations for a minimal number of immunizations as a preferred product characteristic for a new TB vaccine (*20*). However, the current subunit vaccine candidates that are under clinical development are almost exclusively designed as BCG boosting vaccines, each containing one to four antigens shared with BCG (*16*). Therefore, similar to NTM sensitization, the immune responses induced by these vaccines risk inhibiting BCG replication and vaccine efficacy. In contrast, a novel subunit vaccine consisting exclusively of Mtb-specific antigens that are not shared with BCG would ensure non-interference with BCG while simultaneously increasing the overall antigen repertoire with additional Mtb-specific responses. In this way, such a subunit vaccine would ‘complement’ rather than ‘boost’ BCG-induced immunity. Given that live mycobacteria and subunit vaccines induce distinct CD4 T cell responses (*21*), a complementing vaccine would also have improved potential to induce a more diverse population of T cell subsets with distinct effector functions compared to traditional BCG boosting vaccines that will primarily expand pre-existing BCG-imprinted T cells. In addition, BCG has a well-documented adjuvant effect when co-administered with other vaccines in both humans and animals, thus co-administration of BCG with a subunit vaccine could offer the further advantage of potentially increasing immunogenicity of the subunit vaccine (*22–24*).

In this work, we investigate a novel TB vaccine candidate, H107, which combines eight individually protective antigens from Mtb that we confirm are immunogenic in Mtb infected mice and humans, while lacking cross-reactivity to BCG. This eliminates the risk of affecting BCG replication and vaccine take, and enables an unhindered BCG+H107 co-administration regimen. Indeed, a subanalysis of a clinical trial, illustrated that subunit vaccines with BCG-shared antigens do cross-react with BCG *in vivo* and we demonstrate in mice that this cross-reactivity is capable of inhibiting BCG colonization. Interestingly, co-administration of H107 with BCG significantly increases both H107- and BCG-induced immune responses leading to increased BCG efficacy. We use T cell receptor (TCR) sequence analysis to show that H107 vaccination leads to a substantial increase in clonal diversity of the CD4 T cell repertoire over that induced by BCG alone, whereas a traditional BCG boosting vaccine comprising shared Mtb-BCG antigens, H65, has no or little impact. Compared to the H65 vaccine, BCG co-administered with H107 also leads to induction of less differentiated memory CD4 T cells and increased Th17 responses. Collectively, these immunological aspects of BCG+H107 co-administration are associated with markedly increased long-term protection compared to either H107, BCG alone, or BCG co-administered with a subunit vaccine sharing antigens with BCG. This observation supports the development of a co-administration strategy to diversify the vaccine pipeline and increase TB vaccine efficacy in infants, adolescents, and adults.

## RESULTS

### Selection of Mtb-specific vaccine antigens

With the overall goal of designing an Mtb-specific TB vaccine with multiple antigens that do not cross-react with BCG, we identified three initial criteria for selection of vaccine antigens: i) non-immunogenic in the context of BCG vaccination, ii) previously reported immunogenicity in Mtb-infected humans, and iii) induction of protective immune responses in animal models. Current subunit vaccine candidates in the existing TB vaccine pipeline consist of 1-4 antigens (*16*), and through this new candidate vaccine we wanted to expand this number. Since Mtb and BCG share ∼98% of their genes (*25, 26*) we took advantage of the fact that modern BCG strains, such as BCG-Danish and BCG-Pasteur, lack multiple genetic regions of difference (RD) present in Mtb and have additional mutations in gene regulators that control the expression of potential vaccine antigens (*27, 28*). Based on this, we selected eight vaccine targets with confirmed human immune recognition and vaccine potential in animal models (Table 1), but with absent or negligible expression/secretion in either all BCG strains (PPE68, ESAT-6, EspI, EspC, and EspA) or modern BCG strains (MPT64, MPT70, and MPT83) (Table S1). We first confirmed that these antigens were immunogenic in Mtb-infected mice, in which their mean IFN-γ recall responses from splenocytes ranged from 1645 to 174412 pg/ml (Fig. 1A). We next confirmed that there were no immune responses against these proteins after BCG vaccination, as all antigen recall responses were comparable to the media background (Fig. 1B). Finally, we established that the individual antigens were immunogenic by formulating each single recombinant protein with the CAF®01 adjuvant (*29*) and observed immune responses from 839 to 113999 pg/ml IFN-γ in immunized mice (Fig. 1C). Notably, the antigens eliciting the highest immune responses during Mtb infection (ESAT-6 and MPT70) were not the ones giving the highest immune responses after protein vaccination (PPE68, EspI, EspC, and EspA).

**Table 1.**
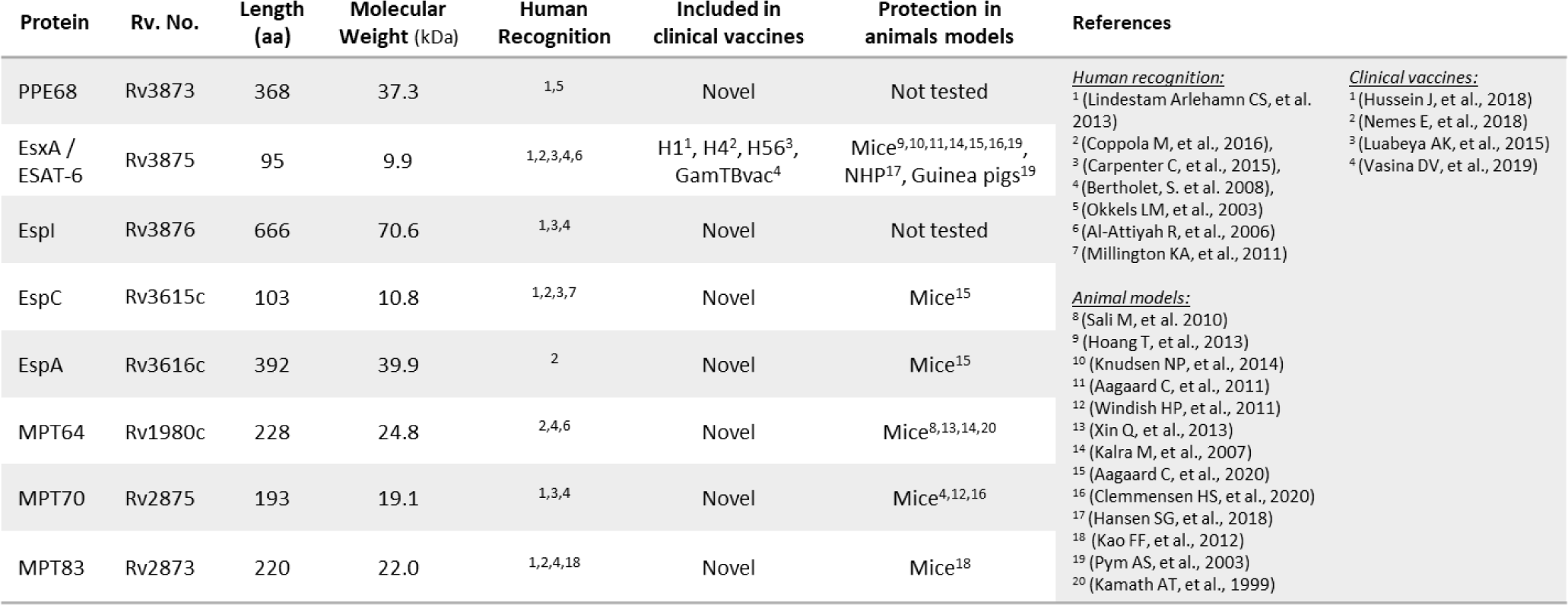
Antigen selection for the H107 vaccine. The H107 vaccine consists of PPE68 (Rv3873), ESAT-6 (Rv3875), EspI (Rv3876, EspC (Rv3615c), EspA (Rv3616c), MPT64 (Rv1980c), MPT70 (Rv2875) and MPT83 (Rv2873). Listed for each of the antigens; Amino acid length, molecular weight of the protein, references to papers showing human recognition, indication of presence in existing vaccine candidates under clinical development as well as protection in animal models.

**Fig. 1.**
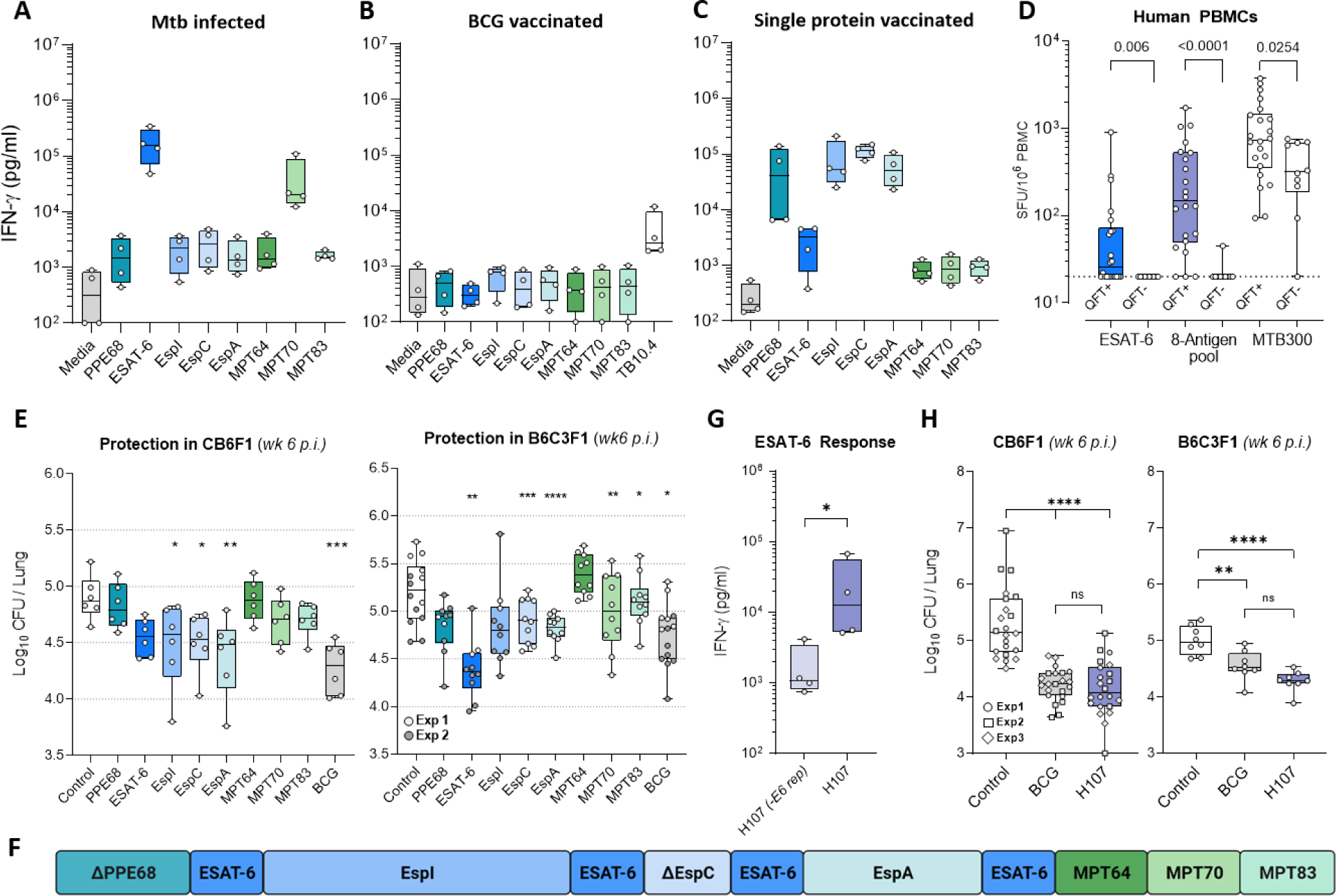
H107 combines immunogenic and protective antigens. Individual antigen responses **(A)** 18 weeks after Mtb challenge, **(B)** eight weeks after vaccination with BCG Danish, or **(C)** two weeks after three subcutaneous (s.c.) immunizations with single recombinant proteins in CAF®01 (n=4) in CB6F1 mice. Splenocytes from BCG-vaccinated mice and Mtb-challenged mice were stimulated *ex vivo* with individual recombinant proteins whereas single-protein vaccinated mice were stimulated with recombinant H107. Levels of IFN-γ in the supernatants after three days of culture were measured by ELISA. **(D)** Magnitude of IFN-γ T cell response in healthy QFT+ (n=22) and QFT- (n=10) subjects against ESAT-6 peptides, the 8-antigen peptide pool, and MTB300. Dotted line indicates the cutoff limit at 20 SFC/106 PBMCs. Two-tailed Mann-Whitney test. **(E)** CB6F1 mice (left) or B6C3F1 mice (right) were immunized three times s.c. with 2 µg individual recombinant proteins in CAF®01, BCG-Danish, or left non-vaccinated, and then challenged with aerosol Mtb Erdman. Lung CFU were determined six weeks post infection (n=6-14). One-Way ANOVA with Dunnett’s multiple comparisons test. **(F)** Antigen design of the H107 fusion protein. Protein modifications can be found in table S2. **(G)** The ESAT-6-specific immune response of splenocytes taken two weeks after three immunizations of CB6F1 mice with 2 µg H107/CAF®01 or H107 that lacks ESAT-6 repeats (-E6 rep)/CAF®01 (n=4). Mann-Whitney test. **(H)** CB6F1 mice (left) and B6C3F1 mice (right) were either vaccinated with H107/CAF®01, BCG-Danish, or left non-vaccinated (Control) and challenged with Mtb Erdman by the aerosol route six weeks post the final vaccination. Lung bacterial burden determined six weeks after challenge (n=8-22). Three independent experiments were performed in CB6F1 mice. One-Way ANOVA with Tukey’s multiple comparisons test. Symbols indicate individual mice or donors. Box plots indicate median, interquartile range, and minimum and maximum values. p-values; *p<0.05, **p<0.01, ***p<0.001, ****p<0.0001, and ns (non-significant).

We finalized the immune characterization by exploring recognition in a cohort of Quantiferon (QFT) positive and negative human subjects from San Diego, USA (Fig. 1D). Peripheral Blood Mononuclear Cells (PBMCs) were obtained from 22 healthy QFT+ and 10 QFT-, US-born individuals. Cells were stimulated with a peptide pool representing all eight antigens and responses were measured by IFN-γ Fluorospot. An ESAT-6 peptide pool and MTB300 (a pool of 300 Mtb-derived human T cell epitopes (*30*)) were included as positive controls. As expected, in the QFT+ cohort significant responses were observed after ESAT-6 stimulation (median 26 SFC/10^6^ PBMCs), where 12/22 QFT+ subjects responded above a cut-off of 20 SFC/10^6^ PBMCs. For the 8-antigen peptide pool, the response magnitude was markedly higher (median 147 SFC/10^6^ PBMCs) and responses were observed in 19/22 QFT+ subjects (Fig. 1D). In the QFT-cohort, only one subject had a response above the cutoff. Interestingly, in contrast to ESAT-6 and the 8-antigen pool, 9/10 QFT-subjects responded to MTB300, albeit with a significantly lower magnitude of response compared to QFT+ subjects (median 318 vs. 739 SFC/10^6^ PBMCs). This suggested cross-reactivity of MTB300 with environmental NTM antigens, while the 8-antigen pool appeared Mtb-specific within this cohort.

We next investigated to what extent immunization with the individual antigens would protect against experimental challenge with Mtb Erdman. To cover multiple different MHC alleles, we performed the experiment in both CB6F1 and B6C3F1 mice. Except for MPT64, all selected antigens induced significant protection in at least one of the two mouse strains (Fig 1E and fig. S1A demonstrating significant protection of PPE68). In CB6F1, the highest bacterial reduction was found for EspA, EspI, ESAT-6, and EspC, whereas ESAT-6 was found to be the single most protective antigen in B6C3F1 followed by EspI, EspA, and EspC. With these results, we designed a fusion protein named H107, that incorporated all the selected antigens (Fig. 1F), including MPT64, as it has well-documented protective capacity in previous studies (*31, 32*). Minor antigen-modifications were introduced in H107 to optimize protein expression and stability, and regions with homology to BCG were removed from PPE68 to ensure Mtb-specificity (Table S2). Also, a C-terminal fragment of Rv3615c was removed to ensure compatibility with the ESAT-6 free diagnostic IGRA, which can then be used as a companion diagnostic (*33*). Since ESAT-6 showed optimal protection in both mouse strains and has been described as a key antigen for sustained vaccine protection both pre- and post-Mtb-exposure (*34–36*), we sought to optimize ESAT-6-specific immune responses. Inspired by experience with immuno-repeats that increased antigenic immunogenicity in a chlamydia subunit vaccine (*37*), we inserted four copies of ESAT-6 into the molecular framework of H107, which led to a significant increase in ESAT-6-specific immunogenicity compared to a version without repeats (Fig. 1G). After immunization with H107 in the CB6F1 strain, responses were strongest for EspI and ESAT-6, followed by responses against EspA and MPT70 (fig. S1B). Finally, we evaluated vaccine efficacy in a short-term aerosol Mtb infection model and found that immunization with H107 in CAF®01 induced protection at least as well as BCG in both CB6F1 and B6C3F1 mice (Fig. 1H).

In summary, we selected eight immunogenic and protective Mtb-specific antigens that were not recognized in context of BCG immunization and that, when combined into a single fusion protein, provided robust protection against pulmonary Mtb infection in mice.

### H107 does not inhibit BCG colonization

Administering TB subunit vaccines while BCG is still replicating comes with the risk that cross-reactive immune responses induced by the subunit vaccine will interact with BCG and inhibit proper colonization and/or exacerbate immune-mediated side effects. This type of interaction was observed during the clinical development of one of the existing TB vaccine candidates, H4:IC31®^1^(*38*) (trial ID: C013-404, NCT02420444) with data reported here for the first time. The H4 protein consists of antigens TB10.4 and Ag85B, which are shared by both BCG and Mtb (*39*). The trial investigated the effects of boosting BCG at different time points using either a two or three-dose regimen (Fig. 2A). In the three-dose regimen, the first dose of H4:IC31® was administered 42 days after BCG prime and led to increased incidences of swelling and erythema at the distal BCG injection site (Fig. 2B, left). However, BCG site reactions were not observed for the two-dose regimen, where the first dose of H4:IC31® was given 98 days post BCG, indicating that the effect was dependent on the time between the administrations (Fig. 2B, middle and right). Overall, this suggests that immune responses induced by the BCG booster vaccine, H4, cross-reacted with BCG at the initial site of injection when administered within a limited window of time. To investigate whether cross-reactive immune responses of BCG booster subunit vaccines have the capacity to inhibit BCG colonization, we used a mouse model. Mice were immunized with CAF®01 mixed with either H107, H4, or the protective subunit hybrid protein H65 (EsxD-EsxC-EsxG-TB10.4-EsxW-EsxV) (*40*), which share zero, two, and six antigens with BCG, respectively. Immunization with the major outer membrane protein (MOMP) from *Chlamydia trachomatis* was included as a control antigen that does not cross-react with BCG. Six weeks after the final immunization, BCG was administered intradermally (i.d.) and analysis of the bacterial load in the spleen 3.5 weeks later showed that both H4- and H65-immunized mice had a reduced BCG load compared to H107- and MOMP-immunized mice (Fig. 2C). Similar results were obtained in a follow-up experiment comparing the effect of H65 and H107 vaccination on BCG colonization in the site-draining lymph nodes (dLNs) (Fig. 2D). Together, these experiments indicated that cross-reactive immune responses induced by traditional antigen-sharing BCG booster vaccines indeed have the capacity to inhibit BCG colonization, whereas H107 did not. As the window for detecting differences in colony forming units (CFU) after i.d. BCG was narrow, given the low number of bacteria in control settings, we further evaluated this in a more sensitive model where BCG was administered intravenously (i.v.) at a higher dose (Fig. 2E). As before, mice were immunized with either H65, H107, or MOMP in CAF®01 and six weeks later subjected to i.v. inoculation with BCG. Here again, H107 or MOMP immunization did not decrease BCG colonization, while H65 vaccination significantly decreased colonization in both lung and spleen (Fig. 2E, left and fig. S2A), an effect that was sustained at least 9 weeks after inoculation. In all studies described so far, and in subsequent figures, the BCG-Danish strain was used. However, given that H107 contains MPT64, MPT70, and MPT83, which are expressed in evolutionarily early BCG strains such as BCG-Russia and BCG-Japan (table S1), we also performed an i.v. infection experiment using BCG-Japan. In agreement with the previous experiment, H65 decreased the BCG-Japan load significantly at week 3.5 post i.v. inoculation, whereas no significant difference was observed 9 weeks after infection in this experiment (Fig. 2E, right and fig. S2A). In contrast, H107 immunization did not affect i.v. BCG-Japan colonization at all time points investigated (Fig. 2E, right). This was despite both H107 (fig. S1B) and BCG-Japan (fig. S2B) inducing detectable MPT70-specific adaptive immune response in this mouse strain.

**Fig. 2.**
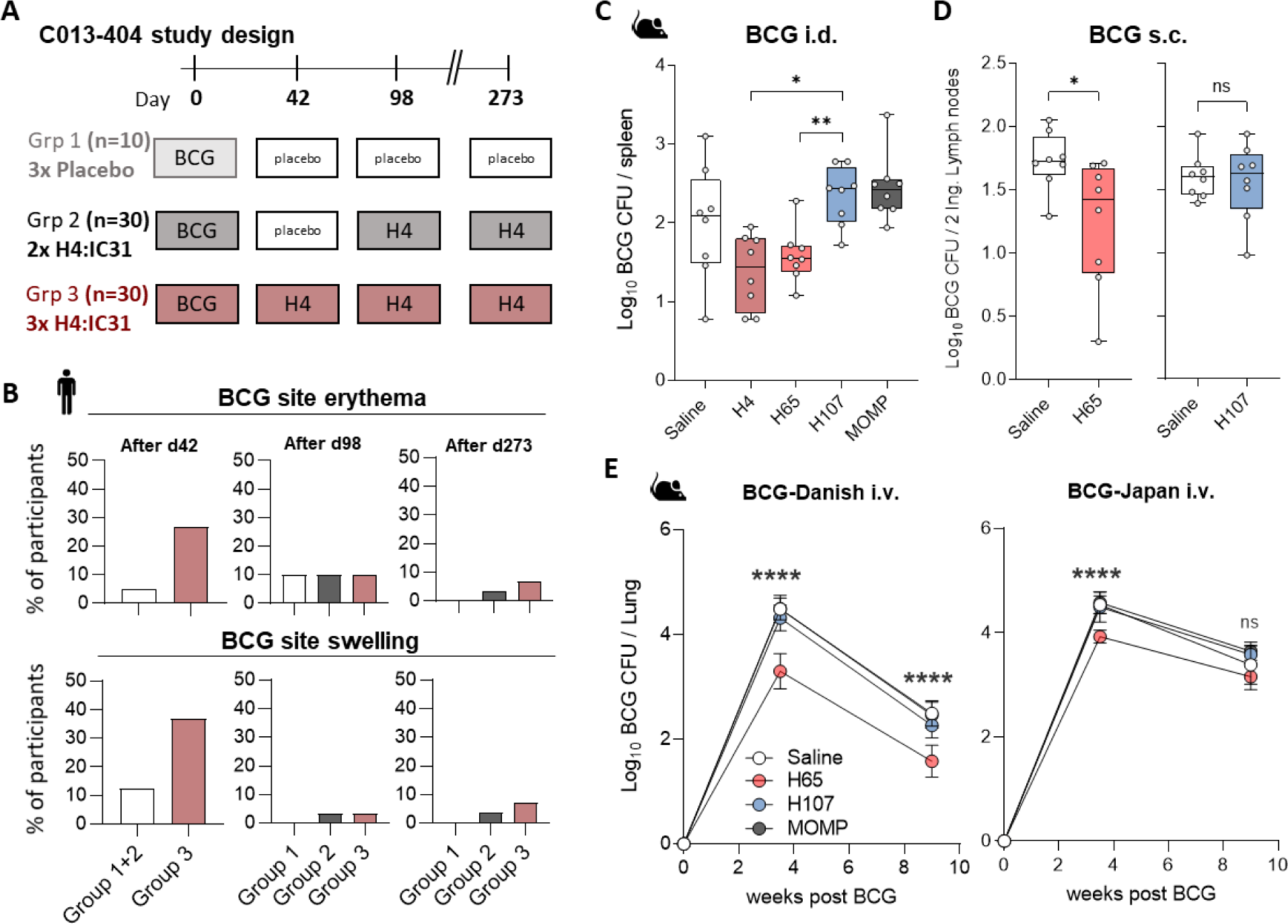
H107 does not induce T cells cross-reactive to BCG. **(A)** Overview of the C013-404 clinical study39. A total of 70 HIV-Negative, TB-Negative, BCG-Naive Adults received BCG-Danish at study day 0 and were randomized on day 42 to either three doses of placebo (Grp. 1, n=10); one dose of placebo followed by two doses of H4:IC31® (Grp. 2, n=30); or three doses of H4:IC31® (Grp. 3, n=30). H4:IC31® or placebo was administered on days 42; 98; 273. **(B)** Reported erythema and swelling at the BCG injection site after each immunization. These adverse events were all mild to moderate in severity. Bars indicate the proportion of individuals with reported events. **(C)** CB6F1 mice were immunized with H4/CAF®01, H107/CAF®01, H65/CAF®01, or MOMP/CAF®01 and inoculated intradermally (i.d.) with BCG and CFUs were determined in the spleens of vaccinated animals 3.5 weeks later (n=8). **(D)** CB6F1 mice were immunized with H65/CAF®01 or H107/CAF®01 and inoculated with BCG subcutaneously (s.c.). BCG CFUs were determined in the inguinal lymph nodes six weeks after inoculation (n=8). **(E)** CB6F1 mice were immunized with H65/CAF®01, H107/CAF®01, and MOMP/CAF®01 and injected intravenously (i.v.) with BCG-Danish (left) or BCG-Japan (right) six weeks after the last immunization. BCG CFUs were enumerated in lungs 3.5 and 9 weeks post BCG challenge (n=8). Data shown as mean± SEM. (C,E) One-way ANOVA with Tukey’s Multiple Comparison test. (D) two-tailed unpaired t-test. p-values; p-values; *p<0.05, **p<0.01, ***p<0.001, ****p<0.0001, and ns (non-significant).

In summary, we demonstrated that BCG booster vaccines containing antigens present in both Mtb and BCG, such as H4:IC31® and H65, have the potential to cross-react with BCG in humans, and that such cross-reactivity can inhibit BCG colonization in mice. Cross-reactivity was not observed for H107, thus enabling a BCG+H107 co-administration regimen.

### BCG and H107 co-administration leads to reciprocal adjuvanticity

Simultaneous co-administration of BCG and subunit vaccines offers several potential advantages, including a reduced number of vaccination visits and the potential for an additional adjuvant effect by BCG on the immunogenicity of the subunit vaccine (*22–24*). Therefore, we next investigated the immunogenicity of a vaccination regimen where BCG was co-administered with the first dose of H107 in CAF®01 (BCG+H107, Fig. 3A). Indeed, compared to a regimen consisting of H107 alone, BCG+H107 co-administration significantly enhanced H107-specific cytokine-producing CD4 T cells measured one week after the final vaccination (Fig. 3B). Though not as prominent as the cellular immune response, the H107-specific antibody responses were also increased by BCG+H107 co-administration, where especially IgG2a and IgG2b titers were increased (fig. S3A). As expected, vaccination with BCG alone did not result in detectable H107-specific cytokine production (Fig. 3B). Targeting BCG to a lymph node distal from the H107-draining lymph node abated the adjuvant effect, in concordance with prior literature describing the need for both vaccines to drain to the same lymph node for a BCG-adjuvant effect (*23*) (fig. S3B). Importantly, the enhancement of the H107 immune response was sustained for up to 16 weeks after the last administration of H107, demonstrating a durable enhancing effect of BCG co-administration on H107 immunogenicity (Fig. 3C). Finally, the increased immunogenicity of H107 in the co-administration regimen was associated with higher levels of pro-inflammatory cytokines in the vaccine-draining lymph node, specifically IFN-γ, IL-1β, KC, and TNFα, indicating increased immune activation (Fig. 3D).

**Fig. 3.**
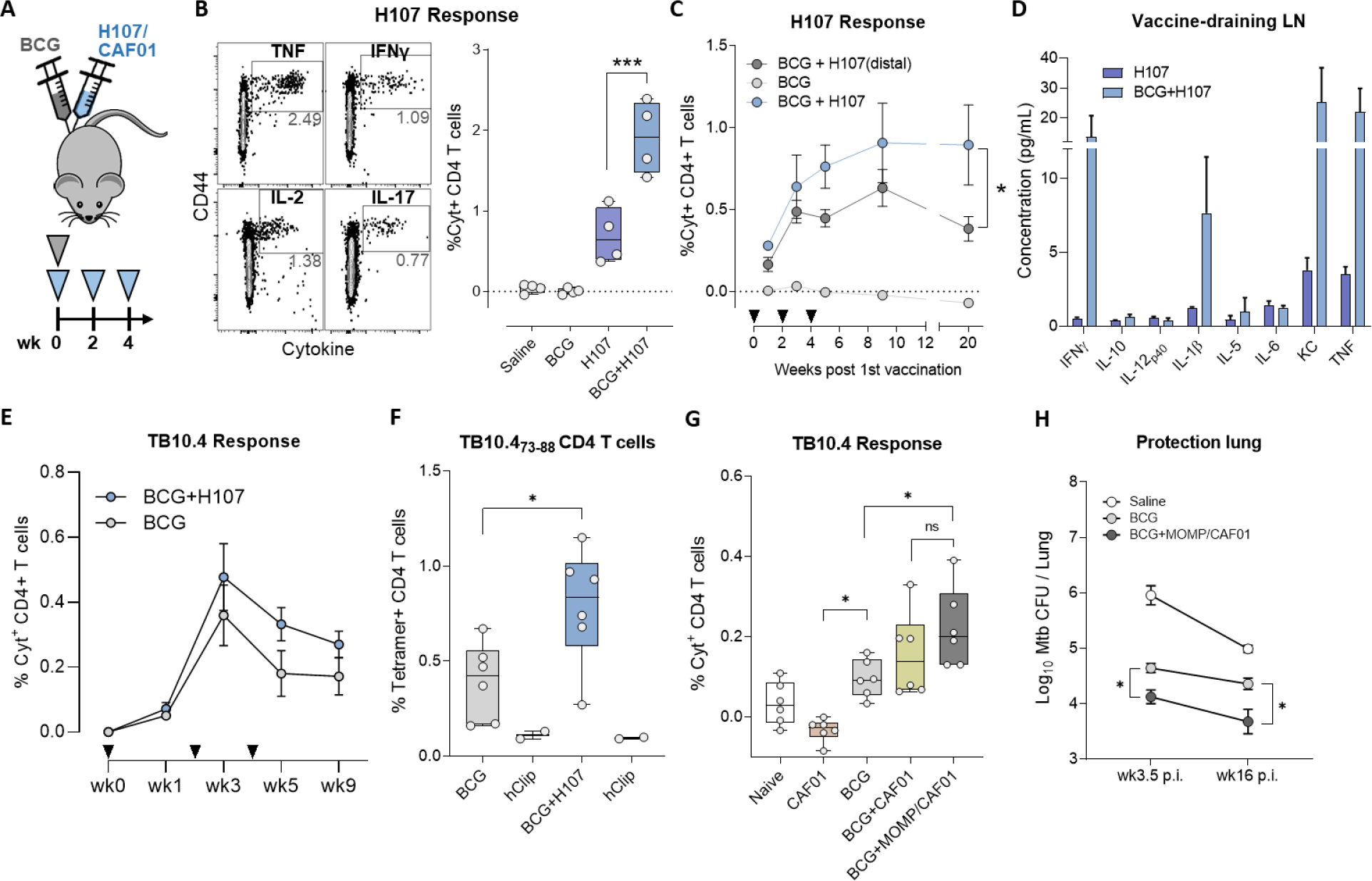
Co-administration of BCG+H107 adjuvants both H107- and BCG-specific immune re-sponses. **(A)** Schematic representation of the BCG+H107 co-administration regimen. Mice were vaccinated once with BCG and H107/CAF®01 s.c. at the base of the tail and boosted twice with H107/CAF®01 s.c., at two-weekly intervals. **(B)** Sample contour plots of gating (left) of ICS analysis to determine the percentage of total cytokine-producing (IFN-γ, IL-2, TNF and/or IL-17A via Boolean OR gating) CD44^high^ CD4 T cells after *ex vivo* restimulation of splenocytes with H107 protein one week post final H107 vaccination and cumulative data for each vaccine group (right). **(C)** H107-specific responses in the spleen over a time course after BCG+H107 co-administration at sites either draining to the same or distal lymph nodes. Triangles indicate vaccination events. **(D)** Cytokine levels in supernatants taken after homogenization of vaccine-draining lymph nodes one week post final H107 vaccination. **(E)** Percentage of cytokine-producing (IFN-γ, IL-2, TNF, and/or IL-17A) CD44^high^ CD4 T cells after restimulating splenocytes with recombinant TB10.4 protein in BCG and BCG+H107 vaccinated mice (n=4, mean ± SEM). Triangles indicate vaccination events. **(F)** TB10.4-specific CD4 T cells in the spleen as determined by I-A^d^:TB10.4_73-88_ tetramer binding one week post final H107 vaccination (n=6). ^I^-Ad:hCLIP_103-117_ negative controls included in duplicates for each group (sample pooled from six animals). Two-tailed unpaired t-test. **(G)** TB10.4-specific T cells one week post final MOMP vaccination measured by ICS as in (E). Mice were vaccinated with buffer only (naïve), CAF®01 adjuvant alone, BCG, BCG+CAF®01 or BCG+MOMP/CAF®01. One-Way ANOVA with Tukey’s Multiple Comparison test. **(H)** Bacterial burden in the lungs 3.5 and 16 weeks post aerosol Mtb infection (p.i.)**. (B,F,G)** Symbols indicate individual mice. Box plots indicate median, interquartile range and minimum and maximum values. **(C-E, H)** Data are plotted as mean±SEM. p-values; *p<0.05, **p<0.01, ***p<0.001, ****p<0.0001, and ns (non-significant).

Having established that BCG acts as a co-adjuvant for H107, we next tracked BCG-specific responses (against TB10.4) to investigate whether H107 immunization would also act as a mutual co-adjuvant for BCG. Intriguingly, compared to BCG alone, BCG+H107 co-administration increased the population of TB10.4-specific CD4 T cells (Fig. 3E), consistent with a parallel significant increase in the frequency of I-A^d^:TB10.4_73-88_ tetramer-positive CD4 T cells detected (Fig. 3F). This reciprocal adjuvant effect appeared to be dependent on the combined presence of both the CAF®01 adjuvant and the protein antigen; co-administration of BCG with unadjuvanted H107 protein did not boost TB10.4-specific cytokine production (fig. S3C), and the increase in TB10.4 response after co-administration with only CAF®01 adjuvant was not significantly increased (Fig. 3G). To study the effect of subunit co-administration on the protective efficacy of BCG in isolation, we co-administered BCG with the unrelated chlamydia-derived protein MOMP formulated with CAF®01. Similar to H107, co-administration of BCG+MOMP significantly increased TB10.4 immune responses (Fig. 3G), which was associated with increased protection compared to BCG alone (Fig. 3H). This effect was maintained into long-term chronic infection (16 weeks) indicating that the synergy of co-administering BCG and subunit vaccines was sustained (Fig. 3H & fig. S3D).

Taken together, these data demonstrate that co-administration of BCG+H107 is feasible and enhances the immunogenicity of both vaccines. Co-administration of BCG with an adjuvanted protein vaccine also enhanced the protection conferred by BCG itself, further arguing for combining BCG and H107 in a co-administration regimen.

### H107 increases clonal diversity compared to a BCG boosting vaccine

We have previously shown that ‘boosting’ pre-existing BCG-induced immunity with Mtb-specific antigens has a greater impact on the CD4 T cell phenotype than traditional BCG boosting where the antigens are shared with BCG (*41*). Based on this, we hypothesized that Mtb-specific vaccines, like H107, drive *de novo* priming of novel T cell clones imprinted by the subunit vaccine, while BCG booster vaccines, such as H65, would largely expand T cell clones initially primed by BCG and thereby have less impact on clonal diversity and T cell phenotype. To test this, we investigated TCR clonal expansion in previously BCG-immunized ‘memory’ mice subsequently ‘boosted’ with either H65 or H107 (Fig. 4A). We confirmed that H65 boosting increased the H65-specific CD4 T cell response and that H107 induced an overall similar magnitude H107-specific response in BCG memory mice (Fig. 4B). Next, we isolated mRNA from purified effector and memory CD4 T cells (CD45RB^low/negative^) from H65 or H107 ‘boosted’ and BCG-memory mice and sequenced the TCR-beta chain locus to identify new clonal clusters induced in the boosted settings. Due to the inherent stochasticity of CDR3 sequences and the resulting lack of overlap in TCR repertoires between individual animals, V-J pairing analysis was used to identify novel T cell “clones”. To normalize for variations in sequencing output, repeated comparisons of equal-sized subsamples of sequences from boosted and BCG-memory samples were performed to identify significantly expanded new clones (Fig. 4B). This analysis showed that H65 boosting resulted in very few new clones, while significantly more new T cell clones were identified after H107 vaccination (Fig. 4C). This suggests that H107 complements the BCG repertoire by priming novel T cell clones, while H65 primarily expands pre-existing BCG-induced T cell populations and therefore has little impact on clonal diversity. In line with this, we observed a greater expansion of the few identified clones in the H65-boost setting (Fig. 4D). Taken together, these findings support that, in contrast to vaccines sharing antigens with BCG, ‘complementing’ vaccines, such H107, promote clonal diversity in the T cell repertoire induced by BCG.

**Fig. 4.**
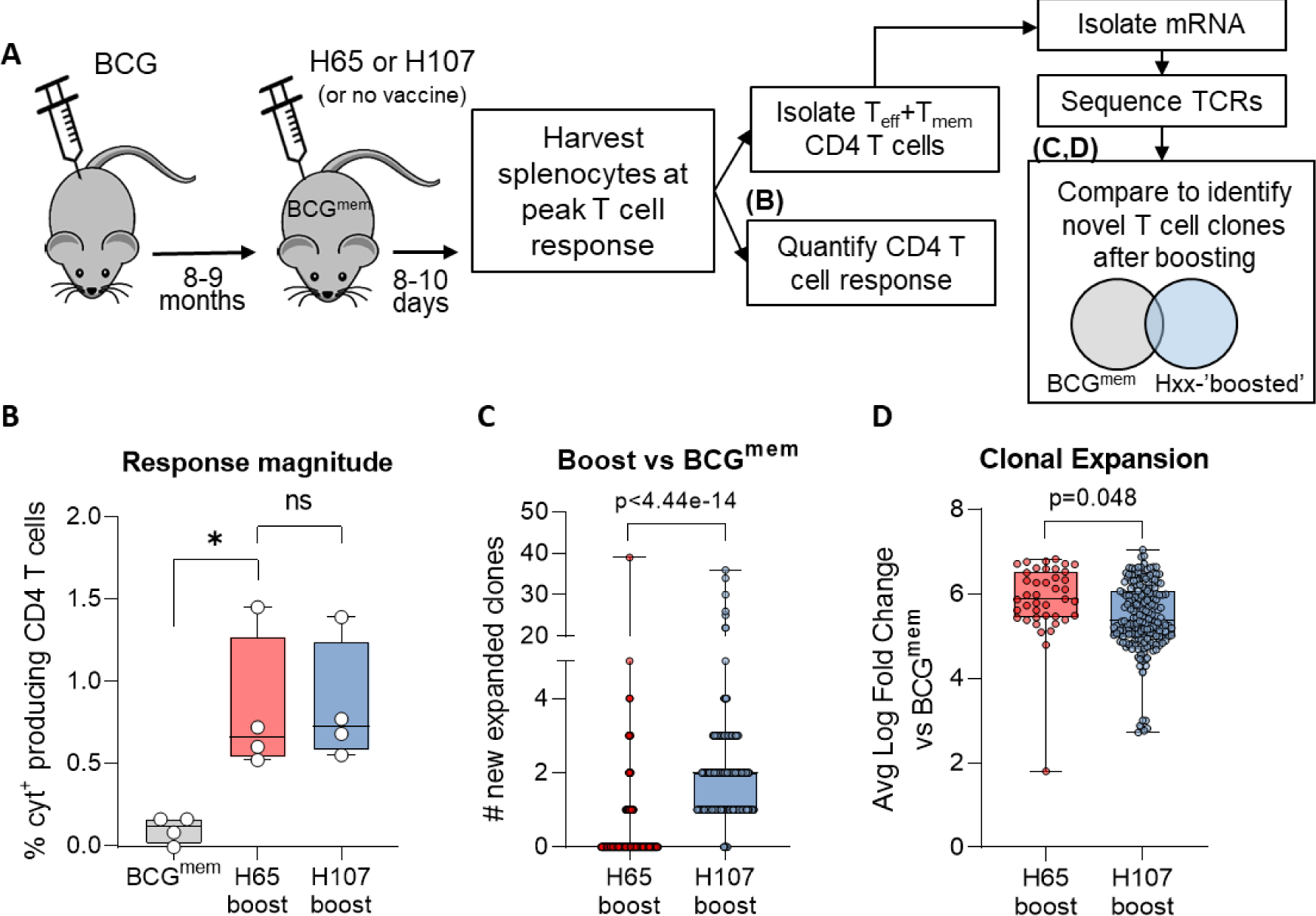
H107 induces novel T cell clones in BCG memory mice. **(A)** Experimental setup in which CB6F1 mice were immunized s.c. with BCG and rested eight-nine months to create memory mice (BCG^mem^). BCG^mem^ mice were then ‘boosted’ by immunizing three times with H65/CAF®01 or H107/CAF®01 at two-week intervals and analyzed eight-ten days after final immunization for CD4 T cell response magnitude via intracellular cytokine staining (ICS) and clonal expansion versus non-boosted BCG^mem^ controls via TCR mRNA sequence analysis. **(B)** Percentage of cytokine-producing (IFN-γ, IL-2, TNF, and/or IL-17A) CD44^high^ CD4 T cells after *ex vivo* restimulation of splenocytes with recombinant H65 or H107 protein eight days after final subunit immunization. Line, median; symbols, individual mice. **(C)** The number of new T cell clones defined by comparison of TCR-β V-J pairings identified in purified T_mem_/T_eff_ CD4 T cells from BCG^mem^ animals with H65- or H107-‘boosted’ mice. **(D)** Average expansion of identified new clones relative to BCG^mem^ control for each subsample comparison in which a significantly expanded novel clonal cluster was identified. **(C,D)** Symbols indicate individual subsampled TCR-β sequence data comparison, p-values determined by Wilcoxon signed rank test with Bonferroni correction.

### BCG co-administered with H107 leads to less differentiated CD4 T cells and increased Th17 responses

We next compared the performance of ‘booster’ versus ‘complementing’ vaccines on CD4 T cell phenotype and functionality in the BCG co-administration setting. As before (Fig. 3A), mice were immunized with BCG-only, or BCG co-administered with either H107 or H65. Immune responses were assessed by antigen stimulation followed by intracellular cytokine staining (ICS) in combination with transcription factor and phenotypic marker expression. Similar to the BCG boosting setting (Fig. 4), co-administration of BCG+H65 increased the H65-specific T cell response over BCG immunization alone, and the magnitude of CD4 T cell responses to the subunit vaccines were similar between BCG+H65 and BCG+H107 immunized animals (Fig. 5A). Principal component analysis (PCA) of the expression of cytokines (IFN-γ/TNF/IL-2 and IL-17), Th1 and Th17 transcription factors (T-bet, RORγT), and surface differentiation and homing markers (KLRG1, CCR7) showed that subunit vaccine-specific CD4 T cell responses in BCG+H65 versus BCG+H107 animals had clearly distinct phenotypes, with differential cytokine expression identified as the major contributor to the variation between groups (Fig. 5B, fig S4A). Further combinatorial expression analysis of Th1 cytokines in BCG-only animals revealed a population of vaccine-specific CD4 T cells with signs of terminal differentiation, including IFN-γ and TNF co-expression (Fig. 5C top), in line with previous studies (*21*).(*42*). In contrast, H107-specific CD4 T cells in BCG+H107 immunized mice displayed a significantly less differentiated phenotype, with reduced IFN-γ expression and increased IL-2 co-expression (Fig. 5C bottom). Notably, H65-specific CD4 T cells in BCG+H65 immunized mice had a phenotype that was intermediate between BCG and BCG+H107 (Fig. 5C middle). To quantitatively compare T cell differentiation between the immunization settings, we utilized a simple functional differentiation score (FDS) that is defined as the ratio of IFN-γ producing T cell subsets divided by the sum of T cell subsets producing other cytokines (IL-2 and/or TNF) but not IFN-γ, as previously described (*34, 41*). As IFN-γ expression is acquired as Th1 cells differentiate, a high FDS score is indicative of a response dominated by differentiated T cells while a low score reveals a less differentiated CD4 T cell population. While BCG+H65 significantly reduced the FDS of vaccine-specific CD4 T cells versus BCG-only primed mice, H107-specific responses in BCG+H107 immunized mice had a still further reduced FDS score, demonstrating that the H107-specific CD4 T cells were less differentiated than those induced by the H65 booster vaccine (Fig. 5D). H107-specific CD4 T cells in BCG+H107 immunized animals also displayed increased IL-17 expression, which is typical of the Th1/Th17-skewing CAF®01 adjuvant, but not s.c. BCG immunization (Fig. 5E) (*43, 44*). In contrast, significantly fewer vaccine-specific CD4 T cells produced IL-17 in animals immunized with BCG+H65 and BCG-alone. Overall, these data support that complementing BCG with H107 more efficiently bypasses BCG-mediated T cell differentiation and Th-imprinting than co-administration of a BCG booster vaccine such as H65.

**Fig. 5.**
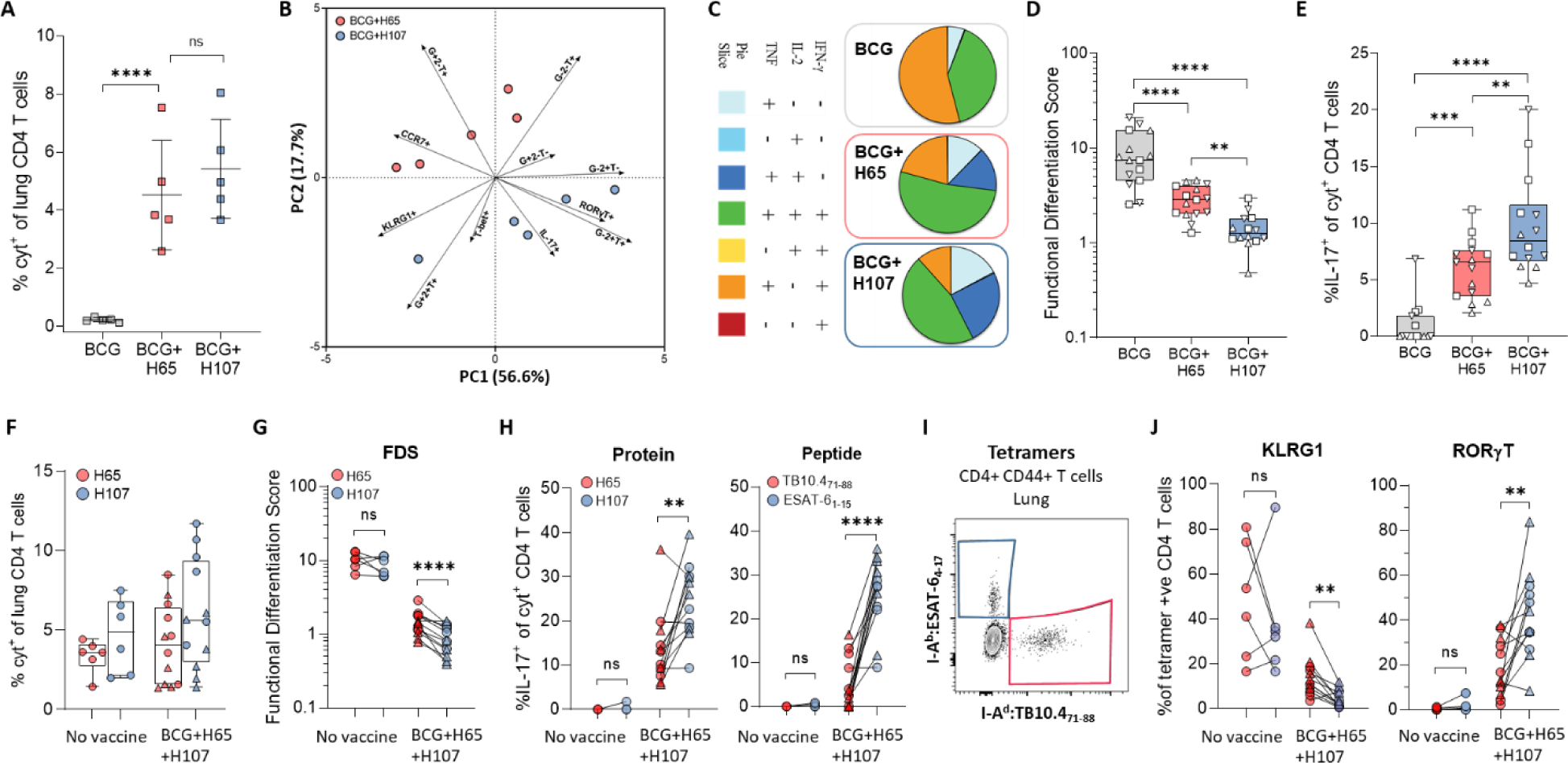
Vaccine-specific T cells induced by BCG+H107 co-administration preferentially acquire a less differentiated profile and Th17 functionality. **(A-D)** CB6F1 mice were immunized s.c. with either BCG, BCG+H65/CAF®01 co-administration, or BCG+H107/CAF®01 co-administration. Five to seven weeks post final immunization, splenocytes of immunized mice were restimulated *ex vivo* with either H65 (BCG, BCG+H65) or H107 (BCG+H107) and assessed by intracellular cytokine staining (ICS). **(A)** Frequency of antigen-specific CD4 T cells producing TNF, IFN-γ, IL-2, or IL-17 after protein restimulation assessed six weeks after the final immunization. **(B)** Principal component analysis (PCA) of percent positive cells for TNF/ IFN-γ/IL-2 (T/G/2), IL-17, RORγT, T-bet, KLRG1, and CCR7 expression of cytokine expressing CD4 T cells from (A). Percentages indicate variance explained by each PC on their respective axis. **(C)** Combinatorial Boolean gating analysis of TNFα/ IFN-γ/IL-2 expression of CD4 T cells from mice analyzed independently five-seven weeks post immunization and depicted as pies indicating the average proportion of antigen-specific T cells with each combination of cytokine expression subset indicated. **(D)** Functional differentiation score (FDS) defined as the ratio of [sum of all IFN-γ producing subsets] : [sum of subsets producing other cytokines (*i.e.* IL-2/TNFα), but not IFN-γ] from mice analyzed five (Δ), six (□), and seven (▽) weeks post immunization. **(E)** Percentage of antigen-specific CD4 T cells expressing IL-17 after *ex vivo* antigen restimulation from mice analyzed independently at five (Δ), six (□), and seven (▽) weeks post immunization. **(F-J)** CB6F1 mice were immunized s.c. by co-administration of BCG and H65+H107 simultaneously in CAF®01 (BCG+H65+H107) or left unimmunized (No Vacc), rested for six weeks, and infected with aerosolized Mtb Erdman. Lung mononuclear cells were isolated at 27 (Δ) and 33 (○) days after Mtb infection and assessed by**(F-H)** *ex vivo* restimulated with H65, H107, TB10.4_71-88_, or ESAT-6_1-15_ as indicated, followed by ICS or **(I, J)** class II MHC tetramer staining. **(F)** Frequency of antigen-specific lung CD4 T cells producing TNF, IFN-γ, IL-2, or IL-17 after protein restimulation, calculated after subtraction of background (media-only stimulated cells). **(G)** Functional differentiation score of cells from (E). **(H)** The proportion of antigen-specific lung CD4 T cells expressing IL-17 after (left) whole vaccine protein stimulation or (right) individual antigenic peptide epitope stimulation. **(I)** A representative contour plot from a BCG+H65+H107 animal 27 days post Mtb infection showing identification of antigen-specific lung CD44^high^ CD4 T cells by I-A^d^:TB10.4_73-88_ or I-A^b^:ESAT-6_4-17_ tetramer binding populations subsequently assessed for **(J)** proportion expressing surface KLRG1 (left) or intracellular RORγT (right) at 27 (Δ) and 33 (○) days post Mtb infection. **(A-J)** Symbols indicate individual animals. Line, mean. Box plots indicate median, IQR, minimum and maximum values. p-values; *p<0.05, **p<0.01, ***p<0.001, ****p<0.0001, and ns (non-significant) based one-way ANOVA with Tukey’s post-test. **(A,D,E)** and paired t-test **(G, H, J)**. Pie slices represent the average relative proportion of each T cell subset of normalized values from individual animals.

Next, we evaluated whether the phenotypic differences of vaccine-specific CD4 T cells were sustained during pulmonary Mtb infection. As the characteristics of T cells are highly influenced by their specific environment, it was important to compare H65- and H107-specific T cells within the same animal with the same bacterial burden. To achieve this, mice were immunized simultaneously with both H65 and H107 in CAF®01, along with BCG co-administration (BCG+H65+H107). In accordance with the results after BCG co-administration with H65 or H107 individually (Fig 5A), BCG+H65+H107 immunized animals had similar H65- and H107-specific CD4 T cell responses post immunization (fig. S4B). After Mtb infection, responses to H65 and H107 were also comparable in immunized as well as non-immunized mice at the site of Mtb infection (Fig. 5F). As expected, Mtb infection-driven lung CD4 T cells had a high FDS in non-immunized mice, where the score was similar between cells specific for both H65 or H107 (Fig. 5G). In contrast, in BCG+H65+H107-immunized mice, the H107-specific CD4 T cell response consistently had a lower FDS score than H65 (Fig. 5G). Furthermore, IL-17 production by Mtb-specific CD4 T cells was present only in BCG+H65+H107 immunized mice, with a significantly higher proportion of H107-specific CD4 T cells expressing IL-17 than H65-specific T cells (Fig. 5H, left). The increased IL-17 production in H107-specific CD4 T cells was further confirmed when comparing single antigenic epitopes within H65 and H107 components, TB10.4 and ESAT-6, respectively (Fig. 5H, right). Parallel analysis of antigen-specific lung CD4 T cells identified by class II MHC tetramers (Fig 5I) confirmed similar magnitudes of the immune responses across the immunized groups (fig. S4C) with increased expression of the Th1 differentiation marker KLRG1 and decreased expression of the Th17 transcription factor RORγT in H65 (TB10.4)-specific versus H107 (ESAT-6)-specific lung CD4 T cells (Fig. 5J). Thus, two independent phenotypic analyses support that decreased T cell differentiation and increased Th17-functionality were preferentially maintained by the H107-induced T cells in response to pulmonary Mtb infection.

Overall, these data demonstrated that BCG co-administered with H107 induced less differentiated CD4 T cells and increased Th17 responses compared to H65. Importantly, these differences were sustained in the adaptive response to pulmonary Mtb infection.

### BCG+H107 co-administration induces superior protection against Mtb challenge

Finally, we evaluated the vaccine efficacy of H107/CAF®01 with and without BCG co-administration, and in relation to BCG and BCG+H65 immunization (Fig. 6). Immunized mice were infected with Mtb by the aerosol route and the bacterial burdens were determined in the lungs at 4 and 18 weeks post infection (p.i.). While BCG+H65 only led to minor improvements over BCG-induced protection (-Δlog 0.26±0.07), co-administration of BCG and H107 induced superior protection compared to BCG (-Δlog 1.26±0.09), BCG+H65 (-Δlog 1.00±0.09) and H107 alone (- Δlog 0.70±0.09) four weeks post infection (Fig. 6, left). Compared to control animals that received saline, where 5/8 animals reached the upper limit of detection for bacterial burden, BCG+H107 induced an impressive 2.94±0.09 log bacterial reduction. The same overall pattern was also observed in a repeat experiment with lower aerosol inoculum (fig.S5). Importantly, while the protective capacity of BCG and BCG+H65 waned over time, BCG+H107 remained significantly protective 18 weeks post infection compared to saline (-Δlog 1.82±0.14), BCG+H65 (-Δlog 1.59±0.14) and H107 (-Δlog 0.66±0.14), demonstrating that the additive protective effect of BCG+H107 outlasted the protective longevity of BCG alone (Fig. 6, right).

**Fig. 6.**
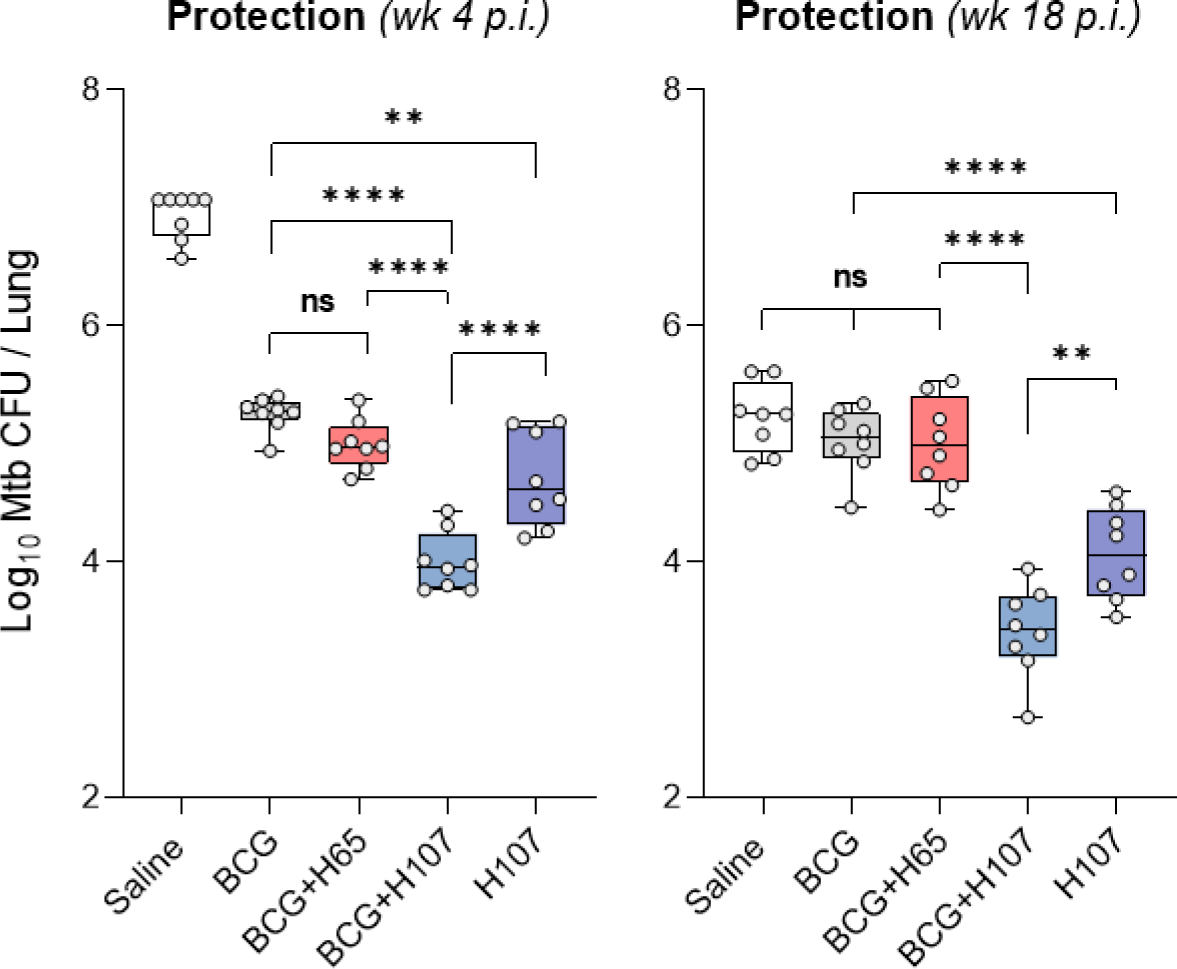
H107 and BCG co-administration induces synergistic protection. CB6F1 mice were immunized once s.c. with BCG, three times s.c. with saline or 1 µg H107/CAF®01, or co-administered BCG with H65/CAF®01 or H107/CAF®01 followed by two subunit boosts(n=8). The bacterial burden was determined in the lungs 4 and 18 weeks post infection (p.i.). Symbols indicate individual mice or donors. Box plots indicate median, interquartile range, and minimum and maximum values. One-Way ANOVA with Tukey’s multiple comparisons test. p-values; **p<0.01, ****p<0.0001, and ns (non-significant).

Collectively, these results showed that H107 induced substantial protection as a stand-alone vaccine (significantly better than BCG), and combining BCG with H107 in a co-administration regimen resulted in further improved long-term protection. This was in contrast to immunization with BCG and BCG+H65, where the protection waned.

## DISCUSSION

Over the past decade, there have been major developments towards a more efficacious TB vaccine (*45*). However, despite the promising results of the M72/AS01E vaccine with 49.7% VE in QFT+ individuals (*14*), more efficacious vaccines are likely needed to end the epidemic (*15*). As the first generation of TB vaccine candidates has moved into more advanced clinical trials, there is a recognized need to diversify the vaccine pipeline to maximize the chance of bringing improved TB vaccines into widespread use (*18, 45, 46*). Such diversity can come in the form of antigenic composition, delivery vector and immune response functionalities. In parallel, recent clinical data demonstrating a potential role for BCG revaccination has also affected how TB vaccine development programs are being moved forward (*47*). Here we describe a novel TB vaccine, H107, that addresses several of these identified shortcomings of first generation TB vaccines while facilitating synergistic interaction with BCG co-administration.

As a specific feature of H107, the fusion protein is selectively composed of eight Mtb-specific antigens that are not expressed or secreted by evolutionary late BCG strains, such as BCG-Danish. It includes the well-described protective Mtb antigen, ESAT-6 (*34–36*), that is repeated four times in the construct to compensate for the low immunogenicity that ESAT-6 has displayed in other subunit vaccines (*34, 48, 49*) (Fig. 1). The remaining seven antigens are novel antigens that have not been tested in clinical trials and significantly expand antigenic coverage compared to the current TB subunit vaccine candidates in the developmental pipeline (*16*). In addition to comprising highly protective antigens (*41*), the Mtb-specific design allows for several novel aspects over other TB vaccine approaches. In contrast to traditional vaccine candidates designed to ‘boost’ BCG, H107 instead ‘complements’ BCG by adding immune responses against additional key protective antigens. This eliminates cross-reactive immune responses with BCG and permits simultaneous co-administration with BCG without negatively impacting BCG colonization (Fig. 2). Importantly, the complementing approach allows the functionalities of the subunit-specific immune response to be regulated by the adjuvant instead of BCG, leading to induction of new Mtb-specific CD4 T cells with improved quality and long-lived immunity (*21, 41*) (Figs. 4,5). In this way, the BCG complementing strategy by H107 is distinct from BCG booster vaccines (e.g. H4, H65) as well as live TB vaccine candidates, where the antigen repertoire is broadened by modifying BCG, or by attenuating Mtb strains via targeted deletions (*36, 50, 51*).

Given the recent promising results in the use of alternative routes of BCG administration against TB (*52, 53*), as well as clinical data showing that revaccination of adolescents/adults with BCG confers protection against Mtb infection (*13*), strategies for combining BCG and subunit vaccine regimens have already been suggested (*19*). In this regard, H107 co-administration could be used as an add-on to further increase protection and compensate for variable BCG efficacy and/or BCG failure in both the existing infant-immunization program as well as potential BCG revaccination programs of the future. In contrast, subunit vaccines sharing antigens with BCG would increase the risk of exacerbating immune-mediated side-effects associated with BCG revaccination and could give rise to lowered BCG vaccine take. Indeed, in the C013-404 BCG boosting study, we observed that administration of H4:IC31® at day 42 post BCG vaccination induced increased swelling and erythema at the BCG injection site, indicating cross-reactivity between the two vaccines. In the mouse model, we confirmed that such cross-reactive immune responses by H4, as well as the BCG boosting vaccine, H65 (*54*), inhibited BCG colonization. This is in line with a previous study showing that immune responses to NTM sensitization can inhibit BCG and decrease protection against experimental Mtb infection (*12*). By design, the antigens included in H107 were not expected to interact with BCG and this was confirmed by a lack of effect of H107 immunization on BCG-Danish colonization in mice (Fig. 2). In the global BCG immunization program, the most commonly used strain is BCG-Danish (or derivatives thereof, such as the Chinese substrain Shanghai D2PB302), but BCG-Russia/Bulgaria and BCG-Japan are also widely used (*55, 56*). We were therefore intrigued to see that H107 immunization also did not affect BCG-Japan colonization in our studies given their potential cross-reactivity inferred by induction of MPT70-specific immune responses by both H107 and BCG-Japan (figs. S1B,S2B). From the data, it is not possible to distinguish whether the lack of impaired i.v. BCG-Japan growth resulted from low immunogenicity of MPT70 following H107 immunization in CB6F1 mice, or a limited antigen availability of MPT70 *in vivo* in the experimental setting. Thus, based on the current study, we conclude that H107 is suitable for co-administration with modern BCG strains, including the VPM1002 vaccine candidate 57 that is in phase II/III trials (NCT03152903, NCT04351685). H107 may also have the potential for co-administration with evolutionary earlier BCG strains, such as BCG-Russia and BCG-Japan, which should be further investigated in future studies.

One of the noteworthy findings of this study was that co-administration of BCG+H107 promotes a significant reciprocal co-adjuvant effect. Consistent with previous reports (*22–24*), we found that BCG adjuvanted the co-administered vaccine, as was reflected in increased H107-specific T cell and antibody responses (Fig. 3, fig. S3). Although the mechanism of such a BCG-adjuvant effect has not been elucidated, we speculate that it is linked to increased inflammation in the draining lymph node, given the observed association with pro-inflammatory cytokines (Fig. 3) (*23*). Interestingly, we also found that subunit immunization had a reciprocal adjuvant effect on BCG immunogenicity and efficacy. This did not depend on shared antigens with BCG, as it was seen with either H107/CAF®01 or MOMP/CAF®01 co-administration. One hypothesis is that increased H107-specific T cell responses in the draining lymph node lead to increased local IL-2 and bystander support of BCG-specific T cell development (*57*). We did not investigate whether H107 co-administration also increases trained non-specific immunity induced by BCG, which could have major implications for the future childhood vaccination program, since BCG is known to decrease overall mortality in early infancy (*58*). Mechanistic investigation of these phenomenon are of high priority in follow-up studies.

CD4 T cells are critical for immune control of Mtb, and T cell differentiation is recognized as a determining factor for their pulmonary protective capacity (*59, 60*). In contrast to live mycobacterial infections like BCG, immunity to adjuvanted protein vaccines is associated with sustained induction of IL-2^+^ less differentiated memory CD4 T cells with superior protective capacity (*21, 61*). More recently, Th17 cells, often induced by mucosal-based immunization regimens, have also been associated with improved CD4 T cell mediated control of Mtb infection (*53*), (*44, 62, 63*). We hypothesized that H107, as a complementing vaccine, would bypass expansion of BCG-imprinted T cells and prime *de novo* adjuvant-imprinted T cells with a quality imprinted by the subunit vaccine itself. This was confirmed by TCR sequencing, where H107 ‘boosting’ of BCG memory was observed to significantly increase clonal diversity over the BCG repertoire whereas H65 boosting had little or no effect. A comparison of the immune responses after co-administration of either BCG+H65 or BCG+H107 showed that BCG+H107 induced a much higher proportion of less-differentiated CD4 T cells that persisted post Mtb challenge and accumulated at the site of infection (Fig. 5). This is in line with our recent work demonstrating that boosting BCG memory with H65 had little influence on T cell quality and protection, whereas boosting with Mtb-specific antigens improved both (*41*). Compared to BCG+H65, we also observed that BCG+H107 induced a significantly increased Th17 response, which is of particular interest given that none of the existing subunit vaccine candidates induce Th17 responses (*46*) despite accumulating evidence of a distinct protective role for Th17 cells (*40, 53, 62, 64-67*). In line with this, we observed that BCG+H107 co-administration led to substantial improvements in protective immunity over both BCG and BCG+H65 (as well as H107 alone) (Fig. 6). The additive effect of H107 and BCG was sustained into chronic infection (week 18) when BCG itself had no impact on the bacterial burden (Fig. 6), and we speculate that the reciprocal adjuvant effect from BCG+H107 co-administration could account for this observation (Fig. 3). Although less than BCG+H107, H107 alone was also significantly more protective than BCG (particularly at week 18), demonstrating robust vaccine potential of H107 as a stand-alone vaccine. Importantly, H65 consists of six highly immunodominant antigens (*54*) and even though co-administration of H65 did not substantially add to BCG mediated protection, we cannot rule out that other subunit vaccines with BCG-shared antigens might. This could be investigated with some of the TB vaccine candidates in the existing pipeline for more rapid implementation of a BCG+subunit co-administration regimen.

Overall, H107 has inherent potential as a standalone vaccine, as well as specific utility in co-administration with BCG vaccination in infants or with BCG (re)vaccination in adolescents and adults. The versatility for BCG co-administration offers a potential pathway for reducing the number of visits in a clinical setting, promoting practical adoption in high TB-burden regions, as described in the WHO preferred product characteristics (*68*). In contrast to BCG boosting vaccines that primarily expand BCG primed T cells, we show that immunization with Mtb-specific antigens has the potential to increase clonal diversity and prime new adjuvant-imprinted CD4 T cells with superior functionality and protection. Based on these unique properties, we believe that H107 has strong translational potential to advance a more effective TB vaccine regimen and have initiated GMP manufacturing in preparation for clinical testing.

## Acknowledgments

We thank the C013-404 study team for providing the clinical data on BCG and H4:IC31® and in particular Dereck Tait and Maria Lempicki (IAVI) for constructive discussions. We acknowledge the NIH Tetramer Core Facility for provision of I-A^b^:ESAT-6_4-17_ and I-A^d^:TB10.4_73-88_ and corresponding negative control tetramers I-A^b^:hCLIP and I-A^d^:hCLIP. We acknowledge Ming Lui Olsen, Camilla Haumann Rasmussen, Camilla Myhre Maymann, and Vivi Andersen for excellent technical assistance in the laboratory as well as the staff at the experimental animal facilities at Statens Serum Institut.

## Author contributions

RM, JSW, HSC, PA, and CA conceived and designed the studies. JSW, HSC, HB, RSL, KD, and TL performed the murine studies and analyzed the data. CSLA designed and analyzed the human PBMC study and JM performed the experiments and analyzed the data. RT recruited the participants and performed the clinical evaluations. IR designed and produced the recombinant proteins including quality control and testing. RJM, JSW, and HSC created the initial draft with critical input and revision for intellectual content from KD, PA, and CSLA. All authors approved the final version.

## Data and materials availability

All data are available in the main text or the supplementary materials. The raw data supporting the conclusions of this article will be made available by the authors, without undue reservation.

## MATERIALS AND METHODS

### Human subjects and samples

Blood samples were obtained from the University of California San Diego, Antiviral Research Center Clinic. Ethical approval to carry out this work is maintained through the La Jolla Institute for Immunology Institutional Review Board. All participants provided written informed consent prior to participation in the study.

We recruited 22 QFT+ individuals and 10 TB negative controls (QFT-). QFT status was confirmed by an IFN-γ release assay (QuantiFERON Gold In-Tube, Cellestis). Further, subjects did not have any clinical or radiographic signs of active TB. Venous blood was collected in heparin-containing blood bags or tubes. PBMCs were purified from whole blood or 100 ml of leukapheresis samples by density-gradient centrifugation (Ficoll-Hypaque; Amersham Biosciences) according to the manufacturer’s instructions. PBMCs were cryopreserved in liquid nitrogen suspended in fetal bovine serum (FBS) (Gemini Bio-Products) containing 10% (vol/vol) DMSO (Sigma-Aldrich).

### Fluorospot assay

T cell responses to ESAT-6, 8-antigen pool, and MTB300 (1µg/ml) peptide pools were measured by IFN-γ Fluorospot assay (Mabtech), according to manufacturer’s instructions. Briefly, Immobilon-FL PVDF 96-well plates (Mabtech) were coated overnight at 4°C with mouse anti-human IFN-γ (clone 1-D1K). PBMCs were thawed and plated at a concentration of 200,000 cells per well and stimulated with the respective proteins and peptide pools at 37°C in a humidified CO2 incubator for 22 hours. As a positive control, 10µg/ml phytohemagglutinin (PHA) was used. In order to assess non-specific cytokine production, cells were also stimulated with DMSO at the corresponding concentration present in the peptide pools, or with culture media alone for the recombinant proteins. All conditions were tested in triplicates. After incubation, cells were removed and plates were washed six times with 200µl PBS with 0.05% Tween 20 using an automated plate washer. After washing, anti-IFN-γ (7-B6-1-FS-BAM) diluted in PBS with 0.1% BSA was added to each well, and plates were incubated for 2 hours at room temperature. The plates were washed again and then incubated with diluted fluorophores (anti-BAM-490) for 1 hour at room temperature. After the final wash, plates were incubated with a fluorescence enhancer for 15 minutes. Spots were counted by computer-assisted image analysis (Mabtech IRIS, Mabtech). The responses were considered positive if they met all three criteria (I) the net spot forming cells per 10^6^ PBMC were >20, (ii) the stimulation index (i.e. fold above background) ≥ 2, and (iii) p≤0.05 by Student’s t-test (*30*).

### Subanalysis of the C013-404 clinical trial (NCT02420444)

This phase I trial was conducted from January 2011 to November 2013 as a double-Blind, randomized, placebo-controlled study at the Centre Hospitalier Universitaire Vaudois, Lausanne, Vaud, Switzerland. A total of 70 HIV-Negative, TB-Negative, BCG-Naive Adults who had received 2-8 × 10^5 CFU BCG-Danish at Study Day −42 were randomized on Study Day 42 in to receive 3 doses of placebo (n=10); 1 dose of placebo followed by 2 doses of 50µg H4 in 500 nmol IC31 (n=30); or 3 doses of 50µg H4 in 500 nmol IC31 (n=30) (see Fig. 2A). All 70 subjects received the first and second vaccinations (days 42 and 98) and 67 (95.7%) subjects completed the full study. The number of unsolicited, solicited, and serious adverse events, including injection site reactions, were recorded from the time of the first vaccination and until the end of the study (Day 259). The study was designed to assess both the safety and immunogenicity of BCG and H4:IC31®, but the data reported in this paper are limited to the reported reactions at the BCG injection site (data on file). The complete dataset from the C013-404 study will be made available in a separate publication. The annotation of C013-404 study timeline in altered this manuscript to parallel the mouse study annotation - ‘day −42’ of C013-404 is re-labelled as ‘day 0’ here (see Fig 2).

### Mice

Six-to-eight week old female B6C3F1 (H2^b,k^) and CB6F1 (H2^b,d^) mice were obtained from Envigo (Netherlands). Mice were randomly assigned to cages of eight upon arrival. Before the initiation of experiments, mice had at least one week of acclimatization in the animal facility. During the course of the experiment, mice had access to irradiated Teklad Global 16% Protein Rodent Diet (Envigo, 2916C) and water *ad libitum*. Mice were housed under Biosafety Level (BSL) II or III conditions in individually ventilated cages (Scanbur, Denmark) and had access to nesting material (enviro-dri and soft paper wool; Brogaarden) as well as enrichment (aspen bricks, paper house, corn, seeds, and nuts; Brogaarden).

### Ethics for animal studies

Statens Serum Institut’s Animal Care and Use Committee approved all experimental procedures and protocols. All experiments were conducted in accordance with the regulations put forward by the Danish Ministry of Justice and Animal Protection Committees under license permit no. 2019-15-0201-00309 and in compliance with the European Union Directive 2010/63 EU.

### Recombinant proteins and fusion proteins

All recombinant proteins and fusion proteins used were produced and purified as previously described (*34*). In brief, DNA constructs were codon-optimized for expression in *E. coli*, inserted into the pJ 411 expression vector (ATUM, Menlo Park, CA, US), expressed, and purified from *E. coli* BL21 (DE3) (Agilent, DK). In the current study, the following individual proteins and fusion proteins were produced: PPE68/Rv3873, ESAT-6/Rv3875, EspI/Rv3876, EspC/Rv3615c, EspA/Rv3616c, MPT70/Rv2875, MPT83/Rv2873, MPT64/Rv1980c, TB10.4/Rv0288, and MOMP and fusion proteins H107, H107*(-E6 rep)* (without ESAT-6 repetition), H4, and H65. The H107 fusion protein is composed of eight Mtb antigens, of which ESAT-6 is repeated four times: PPE68-[ESAT-6]-EspI-[ESAT-6]-EspC-[ESAT-6]-EspA-[ESAT-6]-MPT64-MPT70-MPT83 (Fig. 1, table S1). H107 *(-E6 rep)* is similar to H107, but with only one copy of ESAT-6, between PPE68 and EspI (Fig. 1). H4 (Ag85B-TB10.4) and H65 (EsxD-EsxC-EsxG-TB10.4-EsxW-EsxV) are previously described fusion proteins (*13, 54*). MOMP (a chlamydia antigen) (*69*) was used as a control to detect non-Mtb-specific responses. Protein purity was assessed by SDS-page followed by Coomassie staining and was above 95%.

### Peptide pools

ESAT-6 and H107 peptide pools were obtained from JPT Peptide Technologies GmbH. Peptides were designed as 15-mers with 5-10 amino acids in overlap and a purity >80%. The proteins were covered by the following number of peptides: PPE68 (25 peptides), ESAT-6 (17 peptides), EspI (66 peptides), EspC (9 peptides), EspA (37 peptides), MPT64 (39 peptides), MPT70 (31 peptides), MPT83 (36 peptides).

### Immunization regimens

Mice were immunized three times with two-week intervals (week 0, 2, and 4). Dose volumes were 200 µl for subcutaneously (s.c.) at the base of the tail, 100 µl for intravenously (i.v.) immunization, and 50 µl for intradermal (i.d) immunization. Recombinant proteins or fusion proteins were diluted in Tris-HCL buffer + 9% Trehalose (pH 7.2) to a concentration of 0.1 – 50 µg and formulated in Cationic Adjuvant Formulation 1® (CAF®01) composed of (250 µg DDA / 50 µg TDB) (*29*). BCG-Danish (1331) or BCG-Japan (Tokyo 172) were either diluted in Phosphate Buffered Saline (PBS) or mixed with CAF®01 (Fig. 3.G,H) to a concentration of 0.5×10^6^ BCG Colony Forming Units (CFU) (s.c.), 1.0×10^6^ BCG CFU (i.v.) and 5×10^6^ BCG CFU (i.d.). Mice either received a single dose of BCG in the first round of immunizations or as a “challenge” ten-fifteen weeks after the first immunization with fusion proteins. Negative control mice were immunized with Tris-HCL buffer only, Tris-HCL mixed with CAF®01, or left non-vaccinated.

*BCG co-administration*: In the co-administration regimen, 0.5×10^6^ CFU BCG was administered s.c. on day −1, and the first immunization with adjuvanted fusion protein was administered at the same site on day 0 followed by two adjuvanted protein vaccinations given s.c. at two-week intervals at the opposite side of the first injection (2^nd^) and the same site of injection (3^rd^). In a single experiment (Fig. 3c), BCG and H107 were administered at different sites (H107:neck and BCG:base of tail).

### Mycobacterial infections

Ten weeks after the first immunization, mice were challenged with Mtb Erdman (ATCC 35801 / TMC107). Mtb Erdman was cultured in Difco ™ Middlebrook 7H9 (BD) supplemented with 10% BBL ™ Middlebrook ADC Enrichment (BD) for two-three weeks using an orbital shaker (∼110 rpm, 37°C). Bacteria were harvested in log phase and stored at −80°C until use. On the day of the experiment, the bacterial stock was thawed, sonicated for five minutes, thoroughly suspended with a 27G needle, and mixed with PBS to the desired inoculum dose. Using a Biaera exposure system controlled via AeroMP software, mice were challenged by the aerosol route with virulent Mtb Erdman in a dose equivalent to 50-100 CFUs.

### Enumeration of BCG and Mtb in organs

In order to determine vaccine efficacy or cross-reactivity, BCG and Mtb CFU were enumerated in lungs, spleens, and lymph nodes. Left lung lobes or spleens were homogenized in 3 mL MilliQ water containing PANTA™ Antibiotic Mixture (BD, cat.no. #245114) using GentleMACS M-tubes (Miltenyi Biotec). Lymph nodes were forced through 70-µm cell strainers (BD Biosciences) in 1 mL panta solution. Tissue homogenates were serially diluted, plated onto 7H11 plates (BD), and grown for approximately 14 days at 37°C and 5% CO2. CFU data were log-transformed before analyses.

### Preparation of single-cell suspensions

Spleens, lungs, inguinal lymph nodes were aseptically harvested from euthanized mice. Lungs were first homogenized in Gentle MACS tubes C (Miltenyi Biotec), followed by 1 hour collagenase digestion (Sigma Aldrich; C5138) at 37°C, 5% CO2. The lung homogenate, spleens, and lymph nodes were subsequently forced through 70-µm cell strainers (BD) with the plunger from a 3 mL syringe (BD). Cells were washed twice in cold RPMI or PBS followed by 5 minutes centrifugation at 1800 rpm. Cells were finally resuspended in supplemented RPMI media containing 10% fetal calf serum (FCS) as previously described (*34*). Cells were counted using an automatic Nucleocounter (Chemotec) and cell suspensions were adjusted to 2×10^5^ cells/well for ELISA and 1-2×10^6^ cells/well for flow cytometry.

### IFN-γ ELISA and multiplex cytokine assay

Splenocytes were cultured in the presence of 2 µg/mL recombinant proteins or peptide pools (JPT) for 3 days. Supernatants were harvested and analyzed for levels of IFN-γ using enzyme-linked immunosorbent assay (ELISA) as previously described (*34*).

Lymph node supernatants were harvested after organ homogenization into 1ml media and analyzed for the concentrations of IFN-γ, IL-1β, IL-6, IL-5, KC/GRO, IL-10, IL-12p40 and TNF using Meso Scale Discovery (MSD) according to the manufacturer’s instructions. The plates were read using a Sector Imager 2400 system and cytokine concentrations were determined by 4-parameter logistic non-linear regression analysis of the standard curve.

### CD4 T memory/effector TCR analysis for clonal expansion

Memory and effector CD4 T cells were purified from total splenocytes isolated from immunized mice using EasySep^TM^ Mouse Memory CD4+ T Cell Isolation Kit (StemCell Technologies) as per manufacturer instructions, and purity verified by flow cytometry. Isolated cellular samples were immediately stored in Buffer RLT plus (Qiagen) and kept at −80°C until mRNA isolation. mRNA isolation, sequencing, and *in silico* TCR clonal comparisons were performed with full service bioinformatics firm ENPICOM (Amsterdam, The Netherlands) as follows: The RNA was isolated from 14 samples using Qiagen All Prep DNA/RNA Mini Kit, the lysis step was skipped. The quality of isolated RNA was checked on a Bioanalyzer instrument using Agilent RNA 6000 Pico kit, and all samples were above RIN 9,6 (av. 9,9). The RNA was quantified using Quant-iT RiboGreen RNA assay. All procedures were followed according to the manufacturer’s recommendations. Libraries for each sample were prepared using the maximum RNA template amount allowed with the QIAseq Immune Repertoire RNA Library Kit, the kit produces mixed Alpha and Beta chain libraries that were separated by bioinformatical analysis and sequenced with Illumina’s V3 cartridge (2×300 bp). All reads with identical UMI were deduplicated while keeping their multiplicity (as read count per molecule). Reads with the same UMI but different sequences were converted (corrected) into consensus reads by taking into account both the read counts and the quality of the differing nucleotides. CDR3 lengths were checked, as well as alignment scores of V and J genes. Prior to analysis, the clone tables were filtered to exclude any sequences that were not functional (only in-frame and without stop codons retained). Clones with CDR3s of less than six amino acids were also filtered out. A clone was defined by the unique pairing of the V and J gene with which each extracted receptor is annotated. Clone counts were summarized to reflect each clone definition and these were the counts used for downsampling and differential abundance. After filtering, and prior to any statistical analyses, downsampling was performed to normalize the total clone output across samples. For beta sequences, the clone tables were subsampled to a total of 15000 counts (the smallest sample had 17871 total counts). Each clone was sampled with probability proportional to its assigned count. Tables/samples were subsampled 200 times. Clone counts were normalized using trimmed mean normalization, followed by the voom transform implemented within the limma R package (version 3.46.0) (*70*). A final test for differential abundance was performed on the voom-transformed data. Results were controlled at the adjusted p-value level of 0.15. Significantly expanded clones in the condition of interest (for example in H107-boost versus BCG-memory) were the positive log fold change clones that pass our stringency criterion. Robustly significant clones are clones that appear significant in the majority of subsamples described above. Significantly expanded “new” clones are clones that pass the stringency criterion of the test and are not present in the reference condition (i.e. BCG memory).

### Tetramer staining for flow cytometry analyses

Class II MHC Tetramers (I-A^d^:TB10.4_73-88_:, I-A^b^:ESAT-6_4-17_:) conjugated to Bv421/PE and PE and corresponding negative controls (I-A^d^:hCLIP, I-A^b^:hCLIP) were provided by the NIH tetramer core facility (Atlanta, USA). Single-cell suspensions were stained with tetramers diluted 1:50 in FACS buffer (PBS+1%FCS) containing 1:200 Fc-block (anti-CD16/CD32) for 30 minutes at 37°C, 5% CO2. Tetramer staining was followed by surface staining, fixation, and eventually transcription factor staining as described below.

### *Ex vivo* stimulation for flow cytometry ICS analyses

Splenocytes or lung cells were stimulated *ex vivo* with 2 µg/mL recombinant protein or peptide pools (JPT) in the presence of 1 μg/ml anti-CD28 (clone 37.51) and anti-CD49d (clone 9C10-MFR4.B) for 1 hour at 37°C, 5% CO2followed by the addition of Brefeldin A to 10 µg/mL (Sigma Aldrich; B7651-5mg) and 5-6 hours of additional incubation at 37°C, after which the cells were kept at 4°C until staining.

Cells were stained with surface markers diluted in 50% brilliant stain buffer (BD Horizon; 566349) using anti-CD4-BV510, anti-CD3-BV605, anti-CD8-BV421 or PerCP-Cy5.5, anti-CD44-Alx700, and anti-KLRG1-BV711 at 4°C for 20min., as well as anti-CCR7-PE/Cy7 (@37°C for 30 min.). Fixation and permeabilization were performed with the Fixation/Permeabilization Solution Kit (BD Cytofix/Cytoperm) as per manufacturer’s instructions followed by intracellular cytokine staining (ICS) for anti-IFN-γ-PE-Cy7, anti-IL-2-APC or –APC-Cy7, anti-TNF-PE, anti-IL17A-PerCP-Cy5.5 or -BV421, and anti-CD3-BV605 or –BV650, and/or the Foxp3/Transcription Factor Staining Buffer Set (eBioscience™; 00-5523-00) followed by anti-RORγT-PECF5954 and T-bet-eFl660. Fluorescence minus one controls were performed to set boundaries gates for selected markers. Cells were characterized using a BD LSRFortessa and the FSC files were manually gated with FlowJo v10 (Tree Star).

When identifying antigen-specific cell percentages, the medium-only stimulation condition was subtracted from antigen stimulation for individual samples. For Boolean gating analyses, this media background was subtracted from each Boolean gate individually.

### Statistical analyses

All graphical visualizations and statistical tests were done using GraphPad Prism v8 or v9 (PCA only). The significance of difference between two antigen restimulations within the same animal was assessed with a paired t-test and between two groups of animals with a non-paired t-test. One-Way Analysis of Variance (ANOVA) using Dunnett’s multiple comparison test (comparing to control mice only) or Tukey’s Multiple Comparison test (comparing across all groups) was used to evaluate significant differences between more than two vaccine groups. FDS and CFU values were log-transformed before statistical analysis. For the PCA analysis data were scaled to have a mean of zero and a standard deviation of 1. The specific statistical test used is stated in the figure legends. A p-value above 0.05 was considered not significantly different.

## Supplementary Materials

Fig. S1. Vaccine protection of PPE68 and antigen responses after H107 immunization.

Fig. S2. Cross reactivity of H65 and H107 with BCG-Danish and BCG-Japan.

Fig. S3. H107- and BCG-specific immune responses after co-administration.

Fig. S4. Characterization of immune responses after BCG co-administered with H65 or H107.

Fig. S5. Lung bacterial load in vaccinated and Mtb infected mice.

Fig. S2. Title of the second supplementary figure.

Table S1. Antigen expression of BCG substrains.

Table S2. Overview of antigen modifications in H107.

## Footnotes

1 IC31^®^ is a registered trademark of Valneva Austria GmbH

## References

1. “Global tuberculosis report 2020,” (World Health Organization, Geneva, 2020).

2. A. B. Hogan, B. L. Jewell, E. Sherrard-Smith, J. F. Vesga, O. J. Watson, C. Whittaker, A. Hamlet, J. A. Smith, P. Winskill, R. Verity, M. Baguelin, J. A. Lees, L. K. Whittles, K. E. C. Ainslie, S. Bhatt, A. Boonyasiri, N. F. Brazeau, L. Cattarino, L. V. Cooper, H. Coupland, G. Cuomo-Dannenburg, A. Dighe, B. A. Djaafara, C. A. Donnelly, J. W. Eaton, S. L. van Elsland, R. G. FitzJohn, H. Fu, K. A. M. Gaythorpe, W. Green, D. J. Haw, S. Hayes, W. Hinsley, N. Imai, D. J. Laydon, T. D. Mangal, T. A. Mellan, S. Mishra, G. Nedjati-Gilani, K. V. Parag, H. A. Thompson, H. J. T. Unwin, M. A. C. Vollmer, C. E. Walters, H. Wang, Y. Wang, X. Xi, N. M. Ferguson, L. C. Okell, T. S. Churcher, N. Arinaminpathy, A. C. Ghani, P. G. T. Walker, T. B. Hallett, Potential impact of the COVID-19 pandemic on HIV, tuberculosis, and malaria in low-income and middle-income countries: a modelling study. Lancet Glob Health 8, e1132–e1141 (2020).

3. C. Lange, D. Chesov, J. Heyckendorf, C. C. Leung, Z. Udwadia, K. Dheda, Drug-resistant tuberculosis: An update on disease burden, diagnosis and treatment. Respirology 23, 656–673 (2018).

4. P. E. Fine, Variation in protection by BCG: implications of and for heterologous immunity. Lancet 346, 1339–1345 (1995).

5. A. E. Roth, C. S. Benn, H. Ravn, A. Rodrigues, I. M. Lisse, M. Yazdanbakhsh, H. Whittle, P. Aaby, Effect of revaccination with BCG in early childhood on mortality: randomised trial in Guinea-Bissau. BMJ 340, c671 (2010).

6. L. C. Rodrigues, S. M. Pereira, S. S. Cunha, B. Genser, M. Y. Ichihara, S. C. de Brito, M. A. Hijjar, I. Dourado, A. A. Cruz, C. Sant’Anna, A. L. Bierrenbach, M. L. Barreto, Effect of BCG revaccination on incidence of tuberculosis in school-aged children in Brazil: the BCG-REVAC cluster-randomised trial. Lancet 366, 1290–1295 (2005).

7. C. C. Leung, C. M. Tam, S. L. Chan, M. Chan-Yeung, C. K. Chan, K. C. Chang, Efficacy of the BCG revaccination programme in a cohort given BCG vaccination at birth in Hong Kong. Int J Tuberc Lung Dis 5, 717–723 (2001).

8. Randomised controlled trial of single BCG, repeated BCG, or combined BCG and killed Mycobacterium leprae vaccine for prevention of leprosy and tuberculosis in Malawi. Karonga Prevention Trial Group. Lancet 348, 17-24 (1996).

9. H. C. Poyntz, E. Stylianou, K. L. Griffiths, L. Marsay, A. M. Checkley, H. McShane, Non-tuberculous mycobacteria have diverse effects on BCG efficacy against Mycobacterium tuberculosis. Tuberculosis (Edinb) 94, 226–237 (2014).

10. P. Mangtani, I. Abubakar, C. Ariti, R. Beynon, L. Pimpin, P. E. Fine, L. C. Rodrigues, P. G. Smith, M. Lipman, P. F. Whiting, J. A. Sterne, Protection by BCG vaccine against tuberculosis: a systematic review of randomized controlled trials. Clin Infect Dis 58, 470–480 (2014).

11. M. L. Barreto, S. M. Pereira, D. Pilger, A. A. Cruz, S. S. Cunha, C. Sant’Anna, M. Y. Ichihara, B. Genser, L. C. Rodrigues, Evidence of an effect of BCG revaccination on incidence of tuberculosis in school-aged children in Brazil: second report of the BCG-REVAC cluster-randomised trial. Vaccine 29, 4875–4877 (2011).

12. L. Brandt, J. Feino Cunha, A. Weinreich Olsen, B. Chilima, P. Hirsch, R. Appelberg, P. Andersen, Failure of the Mycobacterium bovis BCG vaccine: some species of environmental mycobacteria block multiplication of BCG and induction of protective immunity to tuberculosis. Infection and immunity 70, 672–678 (2002).

13. E. Nemes, H. Geldenhuys, V. Rozot, K. T. Rutkowski, F. Ratangee, N. Bilek, S. Mabwe,L. Makhethe, M. Erasmus, A. Toefy, H. Mulenga, W. A. Hanekom, S. G. Self, L. G. Bekker, R. Ryall, S. Gurunathan, C. A. DiazGranados, P. Andersen, I. Kromann, T. Evans, R. D. Ellis, B. Landry, D. A. Hokey, R. Hopkins, A. M. Ginsberg, T. J. Scriba, M. Hatherill, C. S. Team, Prevention of M. tuberculosis Infection with H4:IC31 Vaccine or BCG Revaccination. The New England journal of medicine 379, 138–149 (2018).

14. D. R. Tait, M. Hatherill, O. Van Der Meeren, A. M. Ginsberg, E. Van Brakel, B. Salaun, T. J. Scriba, E. J. Akite, H. M. Ayles, A. Bollaerts, M. A. Demoitie, A. Diacon, T. G. Evans, P. Gillard, E. Hellstrom, J. C. Innes, M. Lempicki, M. Malahleha, N. Martinson, D. Mesia Vela, M. Muyoyeta, V. Nduba, T. G. Pascal, M. Tameris, F. Thienemann, R. J. Wilkinson, F. Roman, Final Analysis of a Trial of M72/AS01E Vaccine to Prevent Tuberculosis. The New England journal of medicine 381, 2429–2439 (2019).

15. G. M. Knight, U. K. Griffiths, T. Sumner, Y. V. Laurence, A. Gheorghe, A. Vassall, P. Glaziou, R. G. White, Impact and cost-effectiveness of new tuberculosis vaccines in low- and middle-income countries. Proc Natl Acad Sci U S A 111, 15520–15525 (2014).

16. L. K. Schrager, J. Vekemens, N. Drager, D. M. Lewinsohn, O. F. Olesen, The status of tuberculosis vaccine development. Lancet Infect Dis 20, e28–e37 (2020).

17. B. R. Bloom, New Promise for Vaccines against Tuberculosis. The New England journal of medicine 379, 1672–1674 (2018).

18. G. Voss, D. Casimiro, O. Neyrolles, A. Williams, S. H. E. Kaufmann, H. McShane, M. Hatherill, H. A. Fletcher, Progress and challenges in TB vaccine development. F1000Res 7, 199 (2018).

19. P. Andersen, T. J. Scriba, Moving tuberculosis vaccines from theory to practice. Nat Rev Immunol 19, 550–562 (2019).

20. L. K. Schrager, P. Chandrasekaran, B. H. Fritzell, M. Hatherill, P. H. Lambert, H. McShane, N. Tornieporth, J. Vekemans, WHO preferred product characteristics for new vaccines against tuberculosis. Lancet Infect Dis 18, 828–829 (2018).

21. T. Lindenstrom, A. Moguche, M. Damborg, E. M. Agger, K. Urdahl, P. Andersen, T Cells Primed by Live Mycobacteria Versus a Tuberculosis Subunit Vaccine Exhibit Distinct Functional Properties. EBioMedicine 27, 27–39 (2018).

22. J. Leentjens, M. Kox, R. Stokman, J. Gerretsen, D. A. Diavatopoulos, R. van Crevel, G. F. Rimmelzwaan, P. Pickkers, M. G. Netea, BCG Vaccination Enhances the Immunogenicity of Subsequent Influenza Vaccination in Healthy Volunteers: A Randomized, Placebo-Controlled Pilot Study. J Infect Dis 212, 1930–1938 (2015).

23. J. Dietrich, R. Billeskov, T. M. Doherty, P. Andersen, Synergistic effect of bacillus calmette guerin and a tuberculosis subunit vaccine in cationic liposomes: increased immunogenicity and protection. J Immunol 178, 3721–3730 (2007).

24. M. O. Ota, J. Vekemans, S. E. Schlegel-Haueter, K. Fielding, M. Sanneh, M. Kidd, M. J. Newport, P. Aaby, H. Whittle, P. H. Lambert, K. P. McAdam, C. A. Siegrist, A. Marchant, Influence of Mycobacterium bovis bacillus Calmette-Guerin on antibody and cytokine responses to human neonatal vaccination. Journal of immunology 168, 919–925 (2002).

25. M. A. Behr, M. A. Wilson, W. P. Gill, H. Salamon, G. K. Schoolnik, S. Rane, P. M. Small, Comparative genomics of BCG vaccines by whole-genome DNA microarray. Science 284, 1520–1523 (1999).

26. S. T. Cole, R. Brosch, J. Parkhill, T. Garnier, C. Churcher, D. Harris, S. V. Gordon, K. Eiglmeier, S. Gas, C. E. Barry, 3rd, F. Tekaia, K. Badcock, D. Basham, D. Brown, T. Chillingworth, R. Connor, R. Davies, K. Devlin, T. Feltwell, S. Gentles, N. Hamlin, S. Holroyd, T. Hornsby, K. Jagels, A. Krogh, J. McLean, S. Moule, L. Murphy, K. Oliver, J. Osborne, M. A. Quail, M. A. Rajandream, J. Rogers, S. Rutter, K. Seeger, J. Skelton, R. Squares, S. Squares, J. E. Sulston, K. Taylor, S. Whitehead, B. G. Barrell, Deciphering the biology of Mycobacterium tuberculosis from the complete genome sequence. Nature 393, 537–544 (1998).

27. D. Charlet, S. Mostowy, D. Alexander, L. Sit, H. G. Wiker, M. A. Behr, Reduced expression of antigenic proteins MPB70 and MPB83 in Mycobacterium bovis BCG strains due to a start codon mutation in sigK. Mol Microbiol 56, 1302–1313 (2005).

28. S. Mostowy, A. G. Tsolaki, P. M. Small, M. A. Behr, The in vitro evolution of BCG vaccines. Vaccine 21, 4270–4274 (2003).

29. E. M. Agger, I. Rosenkrands, J. Hansen, K. Brahimi, B. S. Vandahl, C. Aagaard, K. Werninghaus, C. Kirschning, R. Lang, D. Christensen, M. Theisen, F. Follmann, P. Andersen, Cationic liposomes formulated with synthetic mycobacterial cordfactor (CAF01): a versatile adjuvant for vaccines with different immunological requirements. PloS one 3, e3116 (2008).

30. C. S. Lindestam Arlehamn, D. M. McKinney, C. Carpenter, S. Paul, V. Rozot, E. Makgotlho, Y. Gregg, M. van Rooyen, J. D. Ernst, M. Hatherill, W. A. Hanekom, B. Peters, T. J. Scriba, A. Sette, A Quantitative Analysis of Complexity of Human Pathogen-Specific CD4 T Cell Responses in Healthy M. tuberculosis Infected South Africans. PLoS Pathog 12, e1005760 (2016).

31. M. Sali, G. Di Sante, A. Cascioferro, A. Zumbo, C. Nicolo, V. Dona, S. Rocca, A. Procoli, M. Morandi, F. Ria, G. Palu, G. Fadda, R. Manganelli, G. Delogu, Surface expression of MPT64 as a fusion with the PE domain of PE_PGRS33 enhances Mycobacterium bovis BCG protective activity against Mycobacterium tuberculosis in mice. Infection and immunity 78, 5202–5213 (2010).

32. A. T. Kamath, C. G. Feng, M. Macdonald, H. Briscoe, W. J. Britton, Differential protective efficacy of DNA vaccines expressing secreted proteins of Mycobacterium tuberculosis. Infection and immunity 67, 1702–1707 (1999).

33. M. Ruhwald, L. de Thurah, D. Kuchaka, M. R. Zaher, A. M. Salman, A. R. Abdel-Ghaffar, F. A. Shoukry, S. W. Michelsen, B. Soborg, T. Blauenfeldt, S. Mpagama, S. T. Hoff, E. M. Agger, I. Rosenkrands, C. Aagard, G. Kibiki, N. El-Sheikh, P. Andersen, Introducing the ESAT-6 free IGRA, a companion diagnostic for TB vaccines based on ESAT-6. Sci Rep 7, 45969 (2017).

34. H. S. Clemmensen, N. P. H. Knudsen, R. Billeskov, I. Rosenkrands, G. Jungersen, C. Aagaard, P. Andersen, R. Mortensen, Rescuing ESAT-6 Specific CD4 T Cells From Terminal Differentiation Is Critical for Long-Term Control of Murine Mtb Infection. Front Immunol 11, 585359 (2020).

35. T. Hoang, C. Aagaard, J. Dietrich, J. P. Cassidy, G. Dolganov, G. K. Schoolnik, C. V. Lundberg, E. M. Agger, P. Andersen, ESAT-6 (EsxA) and TB10.4 (EsxH) based vaccines for pre- and post-exposure tuberculosis vaccination. PLoS One 8, e80579 (2013).

36. A. S. Pym, P. Brodin, L. Majlessi, R. Brosch, C. Demangel, A. Williams, K. E. Griffiths, G. Marchal, C. Leclerc, S. T. Cole, Recombinant BCG exporting ESAT-6 confers enhanced protection against tuberculosis. Nat Med 9, 533–539 (2003).

37. A. W. Olsen, F. Follmann, K. Erneholm, I. Rosenkrands, P. Andersen, Protection Against Chlamydia trachomatis Infection and Upper Genital Tract Pathological Changes by Vaccine-Promoted Neutralizing Antibodies Directed to the VD4 of the Major Outer Membrane Protein. The Journal of infectious diseases 212, 978–989 (2015).

38. H. Geldenhuys, H. Mearns, D. J. Miles, M. Tameris, D. Hokey, Z. Shi, S. Bennett, P. Andersen, I. Kromann, S. T. Hoff, W. A. Hanekom, H. Mahomed, M. Hatherill, T. J. Scriba, H. I. T. S. Group, M. van Rooyen, J. Bruce McClain, R. Ryall, G. de Bruyn, H. I. T. S. Groupa, The tuberculosis vaccine H4:IC31 is safe and induces a persistent polyfunctional CD4 T cell response in South African adults: A randomized controlled trial. Vaccine 33, 3592–3599 (2015).

39. R. Billeskov, T. T. Elvang, P. L. Andersen, J. Dietrich, The HyVac4 subunit vaccine efficiently boosts BCG-primed anti-mycobacterial protective immunity. PLoS One 7, e39909 (2012).

40. N. P. Knudsen, A. Olsen, C. Buonsanti, F. Follmann, Y. Zhang, R. N. Coler, C. B. Fox, Meinke, U. D’Oro, D. Casini, A. Bonci, R. Billeskov, E. De Gregorio, R. Rappuoli, A. M. Harandi, P. Andersen, E. M. Agger, Different human vaccine adjuvants promote distinct antigen-independent immunological signatures tailored to different pathogens. Sci Rep 6, 19570 (2016).

41. C. Aagaard, N. P. H. Knudsen, I. Sohn, A. A. Izzo, H. Kim, E. H. Kristiansen, T. Lindenstrom, E. M. Agger, M. Rasmussen, S. J. Shin, I. Rosenkrands, P. Andersen, R. Mortensen, Immunization with Mycobacterium tuberculosis-Specific Antigens Bypasses T Cell Differentiation from Prior Bacillus Calmette-Guerin Vaccination and Improves Protection in Mice. Journal of immunology 205, 2146–2155 (2020).

42. C. Aagaard, N. P. H. Knudsen, I. Sohn, A. A. Izzo, H. Kim, E. H. Kristiansen, T. Lindenstrøm, E. M. Agger, M. Rasmussen, S. J. Shin, Immunization with Mycobacterium tuberculosis–Specific Antigens Bypasses T Cell Differentiation from Prior Bacillus Calmette–Guérin Vaccination and Improves Protection in Mice. The Journal of Immunology 205, 2146–2155 (2020).

43. T. Lindenstrom, J. Woodworth, J. Dietrich, C. Aagaard, P. Andersen, E. M. Agger, Vaccine-induced th17 cells are maintained long-term postvaccination as a distinct and phenotypically stable memory subset. Infect Immun 80, 3533–3544 (2012).

44. J. S. Woodworth, D. Christensen, J. P. Cassidy, E. M. Agger, R. Mortensen, P. Andersen, Mucosal boosting of H56:CAF01 immunization promotes lung-localized T cells and an accelerated pulmonary response to Mycobacterium tuberculosis infection without enhancing vaccine protection. Mucosal Immunol 12, 816–826 (2019).

45. M. Hatherill, R. G. White, T. R. Hawn, Clinical Development of New TB Vaccines: Recent Advances and Next Steps. Front Microbiol 10, 3154 (2019).

46. M. J. Rodo, V. Rozot, E. Nemes, O. Dintwe, M. Hatherill, F. Little, T. J. Scriba, A comparison of antigen-specific T cell responses induced by six novel tuberculosis vaccine candidates. PLoS Pathog 15, e1007643 (2019).

47. E. Nemes, H. Geldenhuys, V. Rozot, K. T. Rutkowski, F. Ratangee, N. Bilek, S. Mabwe, L. Makhethe, M. Erasmus, A. Toefy, H. Mulenga, W. A. Hanekom, S. G. Self, L. G. Bekker, R. Ryall, S. Gurunathan, C. A. DiazGranados, P. Andersen, I. Kromann, T. Evans, R. D. Ellis, B. Landry, D. A. Hokey, R. Hopkins, A. M. Ginsberg, T. J. Scriba, M. Hatherill, Prevention of M. tuberculosis Infection with H4:IC31 Vaccine or BCG Revaccination. The New England journal of medicine 379, 138–149 (2018).

48. L. G. Bekker, O. Dintwe, A. Fiore-Gartland, K. Middelkoop, J. Hutter, A. Williams, A. K. Randhawa, M. Ruhwald, I. Kromann, P. L. Andersen, C. A. DiazGranados, K. T. Rutkowski, D. Tait, M. D. Miner, E. Andersen-Nissen, S. C. De Rosa, K. E. Seaton, G. D. Tomaras, M. J. McElrath, A. Ginsberg, J. G. Kublin, H. A. A.-P. Team, A phase 1b randomized study of the safety and immunological responses to vaccination with H4:IC31, H56:IC31, and BCG revaccination in Mycobacterium tuberculosis-uninfected adolescents in Cape Town, South Africa. EClinicalMedicine 21, 100313 (2020).

49. C. Aagaard, T. Hoang, J. Dietrich, P. J. Cardona, A. Izzo, G. Dolganov, G. K. Schoolnik, J. P. Cassidy, R. Billeskov, P. Andersen, A multistage tuberculosis vaccine that confers efficient protection before and after exposure. Nat Med 17, 189–194 (2011).

50. N. Aguilo, J. Gonzalo-Asensio, S. Alvarez-Arguedas, D. Marinova, A. B. Gomez, S. Uranga, R. Spallek, M. Singh, R. Audran, F. Spertini, C. Martin, Reactogenicity to major tuberculosis antigens absent in BCG is linked to improved protection against Mycobacterium tuberculosis. Nature communications 8, 16085 (2017).

51. D. Bottai, W. Frigui, S. Clark, E. Rayner, A. Zelmer, N. Andreu, M. I. de Jonge, G. J. Bancroft, A. Williams, P. Brodin, R. Brosch, Increased protective efficacy of recombinant BCG strains expressing virulence-neutral proteins of the ESX-1 secretion system. Vaccine 33, 2710–2718 (2011).

52. P. A. Darrah, J. J. Zeppa, P. Maiello, J. A. Hackney, M. H. Wadsworth, 2nd, T. K. Hughes, S. Pokkali, P. A. Swanson, 2nd, N. L. Grant, M. A. Rodgers, M. Kamath, C. M. Causgrove, D. J. Laddy, A. Bonavia, D. Casimiro, P. L. Lin, E. Klein, A. G. White, C. A. Scanga, A. K. Shalek, M. Roederer, J. L. Flynn, R. A. Seder, Prevention of tuberculosis in macaques after intravenous BCG immunization. Nature 577, 95–102 (2020).

53. K. Dijkman, C. C. Sombroek, R. A. W. Vervenne, S. O. Hofman, C. Boot, E. J. Remarque, C. H. M. Kocken, T. H. M. Ottenhoff, I. Kondova, M. A. Khayum, K. G. Haanstra, M. P. M. Vierboom, F. A. W. Verreck, Prevention of tuberculosis infection and disease by local BCG in repeatedly exposed rhesus macaques. Nat Med 25, 255–262 (2011).

54. N. P. Knudsen, S. Norskov-Lauritsen, G. M. Dolganov, G. K. Schoolnik, T. Lindenstrom, P. Andersen, E. M. Agger, C. Aagaard, Tuberculosis vaccine with high predicted population coverage and compatibility with modern diagnostics. Proc Natl Acad Sci U S A 111, 1096–1101 (2011).

55. T. Cernuschi, S. Malvolti, E. Nickels, M. Friede, Bacillus Calmette-Guerin (BCG) vaccine: A global assessment of demand and supply balance. Vaccine 36, 498–506 (2011).

56. N. Ritz, N. Curtis, Mapping the global use of different BCG vaccine strains. Tuberculosis (Edinb) 89, 248–251 (2011).

57. A. C. Antoniou, O. M. Sinilnikova, L. McGuffog, S. Healey, H. Nevanlinna, T. Heikkinen, J. Simard, A. B. Spurdle, J. Beesley, X. Chen, C. Kathleen Cuningham Foundation Consortium for Research into Familial Breast, S. L. Neuhausen, Y. C. Ding, F. J. Couch, X. Wang, Z. Fredericksen, P. Peterlongo, B. Peissel, B. Bonanni, A. Viel, L. Bernard, P. Radice, C. I. Szabo, L. Foretova, M. Zikan, K. Claes, M. H. Greene, P. L. Mai, G. Rennert, F. Lejbkowicz, I. L. Andrulis, H. Ozcelik, G. Glendon, Ocgn, A. M. Gerdes, M. Thomassen, L. Sunde, M. A. Caligo, Y. Laitman, T. Kontorovich, S. Cohen, B. Kaufman, E. Dagan, R. G. Baruch, E. Friedman, K. Harbst, G. Barbany-Bustinza, J. Rantala, H. Ehrencrona, P. Karlsson, S. M. Domchek, K. L. Nathanson, A. Osorio, I. Blanco, A. Lasa, J. Benitez, U. Hamann, F. B. Hogervorst, M. A. Rookus, J. M. Collee, P. Devilee, M. J. Ligtenberg, R. B. van der Luijt, C. M. Aalfs, Q. Waisfisz, J. Wijnen, C. E. van Roozendaal, Hebon, S. Peock, M. Cook, D. Frost, C. Oliver, R. Platte, D. G. Evans, F. Lalloo, R. Eeles, L. Izatt, R. Davidson, C. Chu, D. Eccles, T. Cole, S. Hodgson, Embrace, A. K. Godwin, D. Stoppa-Lyonnet, B. Buecher, M. Leone, B. Bressac-de Paillerets, A. Remenieras, O. Caron, G. M. Lenoir, N. Sevenet, M. Longy, S. F. Ferrer, F. Prieur, Gemo, D. Goldgar, A. Miron, E. M. John, S. S. Buys, M. B. Daly, J. L. Hopper, M. B. Terry, Y. Yassin, R. Breast Cancer Family, C. Singer, D. Gschwantler-Kaulich, C. Staudigl, T. Hansen, R. B. Barkardottir, T. Kirchhoff, P. Pal, K. Kosarin, K. Offit, M. Piedmonte, G. C. Rodriguez, K. Wakeley, J. F. Boggess, J. Basil, P. E. Schwartz, S. V. Blank, A. E. Toland, M. Montagna, C. Casella, E. N. Imyanitov, A. Allavena, R. K. Schmutzler, B. Versmold, C. Engel, A. Meindl, N. Ditsch, N. Arnold, D. Niederacher, H. Deissler, B. Fiebig, C. Suttner, I. Schonbuchner, D. Gadzicki, T. Caldes, M. de la Hoya, K. A. Pooley, D. F. Easton, G. Chenevix-Trench, Cimba, Common variants in LSP1, 2q35 and 8q24 and breast cancer risk for BRCA1 and BRCA2 mutation carriers. Human molecular genetics 18, 4442–4456 (2009).

58. J. P. Higgins, K. Soares-Weiser, J. A. Lopez-Lopez, A. Kakourou, K. Chaplin, H. Christensen, N. K. Martin, J. A. Sterne, A. L. Reingold, Association of BCG, DTP, and measles containing vaccines with childhood mortality: systematic review. BMJ 355, i5170 (2016).

59. S. Sakai, K. D. Kauffman, J. M. Schenkel, C. C. McBerry, K. D. Mayer-Barber, D. Masopust, D. L. Barber, Cutting edge: control of Mycobacterium tuberculosis infection by a subset of lung parenchyma-homing CD4 T cells. J Immunol 192, 2965–2969 (2011).

60. M. A. Sallin, S. Sakai, K. D. Kauffman, H. A. Young, J. Zhu, D. L. Barber, Th1 Differentiation Drives the Accumulation of Intravascular, Non-protective CD4 T Cells during Tuberculosis. Cell Rep 18, 3091–3104 (2017).

61. A. O. Moguche, M. Musvosvi, A. Penn-Nicholson, C. R. Plumlee, H. Mearns, H. Geldenhuys, E. Smit, D. Abrahams, V. Rozot, O. Dintwe, S. T. Hoff, I. Kromann, M. Ruhwald, P. Bang, R. P. Larson, S. Shafiani, S. Ma, D. R. Sherman, A. Sette, C. S. Lindestam Arlehamn, D. M. McKinney, H. Maecker, W. A. Hanekom, M. Hatherill, P. Andersen, T. J. Scriba, K. B. Urdahl, Antigen Availability Shapes T Cell Differentiation and Function during Tuberculosis. Cell Host Microbe 21, 695–706 e695 (2017).

62. C. Counoupas, K. C. Ferrell, A. Ashhurst, N. D. Bhattacharyya, G. Nagalingam, E. L. Stewart, C. G. Feng, N. Petrovsky, W. J. Britton, J. A. Triccas, Mucosal delivery of a multistage subunit vaccine promotes development of lung-resident memory T cells and affords interleukin-17-dependent protection against pulmonary tuberculosis. NPJ Vaccines 5, 105 (2020).

63. E. Van Dis, K. M. Sogi, C. S. Rae, K. E. Sivick, N. H. Surh, M. L. Leong, D. B. Kanne, K. Metchette, J. J. Leong, J. R. Bruml, V. Chen, K. Heydari, N. Cadieux, T. Evans, S. M. McWhirter, T. W. Dubensky, Jr., D. A. Portnoy, S. A. Stanley, STING-Activating Adjuvants Elicit a Th17 Immune Response and Protect against Mycobacterium tuberculosis Infection. Cell Rep 23, 1435–1447 (2018).

64. S. A. Khader, G. K. Bell, J. E. Pearl, J. J. Fountain, J. Rangel-Moreno, G. E. Cilley, F. Shen, S. M. Eaton, S. L. Gaffen, S. L. Swain, R. M. Locksley, L. Haynes, T. D. Randall, A. M. Cooper, IL-23 and IL-17 in the establishment of protective pulmonary CD4+ T cell responses after vaccination and during Mycobacterium tuberculosis challenge. Nat Immunol 8, 369–377 (2011).

65. R. Gopal, J. Rangel-Moreno, S. Slight, Y. Lin, H. F. Nawar, B. A. Fallert Junecko, T. A. Reinhart, J. Kolls, T. D. Randall, T. D. Connell, S. A. Khader, Interleukin-17-dependent CXCL13 mediates mucosal vaccine-induced immunity against tuberculosis. Mucosal Immunol 6, 972–984 (2011).

66. L. Monin, K. L. Griffiths, S. Slight, Y. Lin, J. Rangel-Moreno, S. A. Khader, Immune requirements for protective Th17 recall responses to Mycobacterium tuberculosis challenge. Mucosal Immunol 8, 1099–1109 (2011).

67. U. Shanmugasundaram, A. N. Bucsan, S. R. Ganatra, C. Ibegbu, M. Quezada, R. V. Blair, X. Alvarez, V. Velu, D. Kaushal, J. Rengarajan, Pulmonary Mycobacterium tuberculosis control associates with CXCR3- and CCR6-expressing antigen-specific Th1 and Th17 cell recruitment. JCI Insight 5, (2020).

68. “WHO Preferred Product Characteristics for New Tuberculosis Vaccines,” (World Health Organization, Geneva, 2018).

69. J. Hansen, K. T. Jensen, F. Follmann, E. M. Agger, M. Theisen, P. Andersen, Liposome delivery of Chlamydia muridarum major outer membrane protein primes a Th1 response that protects against genital chlamydial infection in a mouse model. J Infect Dis 198, 758–767 (2011).

70. M. E. Ritchie, B. Phipson, D. Wu, Y. Hu, C. W. Law, W. Shi, G. K. Smyth, limma powers differential expression analyses for RNA-sequencing and microarray studies. Nucleic Acids Res 43, e47 (2015).

71. J. Liu, V. Tran, A. S. Leung, D. C. Alexander, B. Zhu, BCG vaccines: their mechanisms of attenuation and impact on safety and protective efficacy. Hum Vaccin 5, 70–78 (2011).

72. L. M. Okkels, I. Brock, F. Follmann, E. M. Agger, S. M. Arend, T. H. Ottenhoff, F. Oftung, I. Rosenkrands, P. Andersen, PPE protein (Rv3873) from DNA segment RD1 of Mycobacterium tuberculosis: strong recognition of both specific T-cell epitopes and epitopes conserved within the PPE family. Infect Immun 71, 6116–6123 (2011).

73. M. Zhang, J. M. Chen, C. Sala, J. Rybniker, N. Dhar, S. T. Cole, EspI regulates the ESX-1 secretion system in response to ATP levels in Mycobacterium tuberculosis. Mol Microbiol 93, 1057–1065 (2011).

