## Supplemental Materials for "A novel Mycobacterium tuberculosis-specific subunit vaccine provides synergistic immunity upon co-administration with Bacillus Calmette-Guérin"

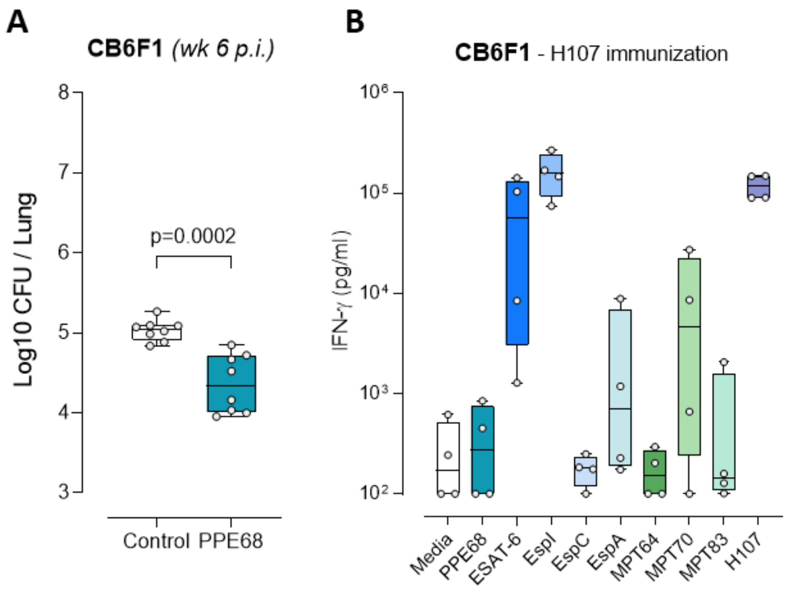


**Fig. S1. Vaccine protection of PPE68 and antigen responses after H107 immunization.**

**(A)** Bacterial numbers were determined in the lungs of control (non-vaccinated) and PPE68-vaccinated mice six weeks post aerosol Mtb challenge (n=8). Two-tailed unpaired t-test. **(B)** CB6F1 mice were vaccinated with H107 three times s.c. and splenocytes were harvested two weeks after the third vaccination. Splenocytes were restimulated *ex vivo* with medium, individual recombinant antigens, or recombinant H107 for three days. The levels of IFN-γ (pg/mL) were measured in the culture supernatant (n=4). Symbols indicate individual mice. Box plots indicate median, interquartile range, and minimum and maximum values.

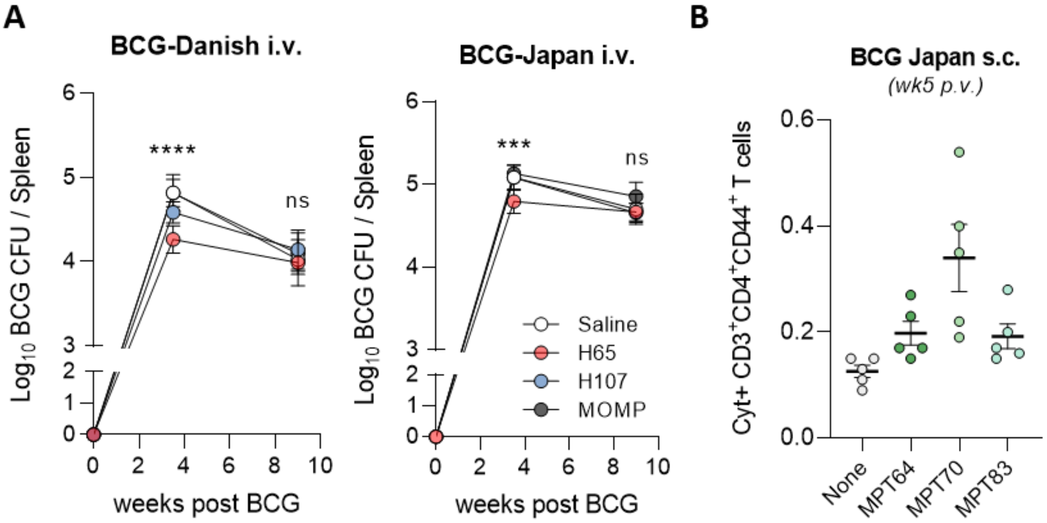


**Fig. S2. Cross reactivity of H65 and H107 with BCG-Danish and BCG-Japan**

**(A)** Inhibition of BCG-Danish and BCG-Japan colonization in the spleen. CB6F1 mice were immunized with H65/CAF®01, H107/CAF®01, or MOMP/CAF®01 and then injected with BCG intravenously (i.v.) six weeks after the last immunization. BCG CFUs were enumerated in the spleen 3.5 and 9 weeks post BCG inoculation (n=8). One-way ANOVA with Tukey's Multiple Comparison test. p-values; ***(p<0.001), ****(p<0.0001) and ns (non-significant). **(B)** MPT70, MPT83, and MPT64-specific immune responses induced by BCG-Japan. CB6F1 mice were vaccinated with BCG-Japan s.c. and the immune responses were assessed five weeks post vaccination (n=4).

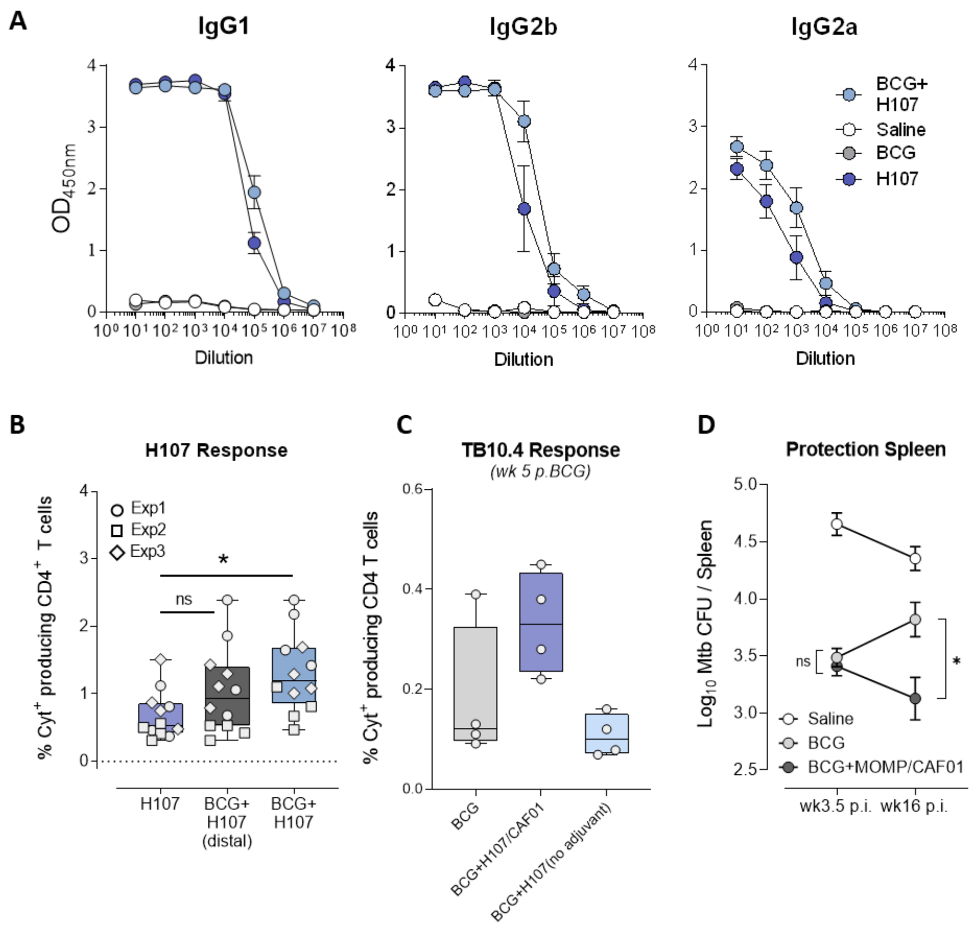


**Fig. S3. H107- and BCG-specific immune responses after co-administration**

**(A)** CB6F1 mice were vaccinated with saline, BCG-Danish, H107/CAF®01, or a co-administration regimen of BCG+H107. One week after final H107 immunization, H107-specific antibodies (IgG1, IgG2b, and IgG2a) measured in blood plasma by ELISA (n=4). **(B)** Comparison of cytokine production after BCG+H107 were co-administered such as to drain to the same lymph node or different, distal lymph nodes. The percentage of total cytokine-producing (IFN-γ, IL-2, TNF and/or IL-17A via Boolean OR gating) CD44$^{high}$ CD4 T cells after *ex vivo* restimulation of splenocytes with H107 protein one week post final H107 vaccination are shown. Data compiled from three independent experiments, as indicated. **(C)** CB6F1 mice were vaccinated with BCG-Danish or a co-administration regimen of BCG+H107 with or without adjuvant CAF®01. Five weeks after BCG (one week after the final H107 vaccination), the percentage of total cytokine-producing CD44$^{high}$ CD4 T cells after *ex vivo* restimulation of splenocytes with TB10.4 protein (n=4). **(D)** CB6F1 mice were vaccinated s.c. with saline, BCG-Danish, or BCG+MOMP/CAF®01 followed by two MOMP/CAF®01 boosts. Mice were rested for 6 weeks and then challenged with aerosolized Mtb Erdman (n=7-8). The bacterial burden was accessed in the spleen 3.5 and 16 weeks post Mtb infection (p.i.). One-Way ANOVA with Tukey's multiple comparisons test. **(A, D)** Data plotted as average mean ± SEM of individual mice. **(B, C)** Symbols indicate individual mice. Box plots indicate median, interquartile range, and minimum and maximum values.  p-values; *($p<0.05$), and ns (non-significant).

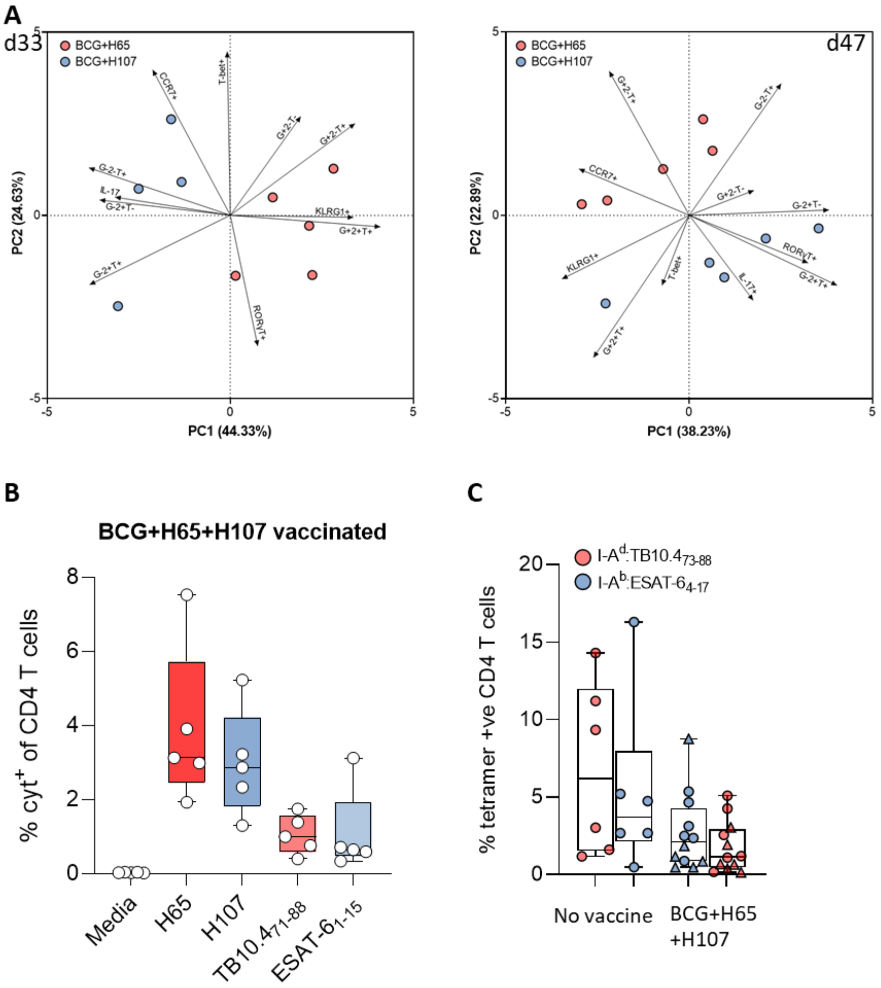


**Fig. S4. Characterization of immune responses after BCG co-administered with H65 or H107**

**(A)** CB6F1 mice were immunized s.c. with either BCG+H65/CAF®01 or BCG+H107/CAF®01. Five (left) and seven (right) weeks post final immunization, splenocytes of immunized mice were restimulated ex vivo with H65 (BCG+H65, red) or H107 (BCG+H107, blue), and the cytokine positive cells determined by ICS. Principal component analysis (PCA) of percent positive CD4 T cells for TNF/ IFN-γ/IL-2, IL-17, RORγT, T-bet, KLRG1, and CCR7 expression of samples. Percentages indicate variance explained by each PC on their respective axis. **(B)** CB6F1 mice were immunized s.c. by co-administration of BCG+ simultaneous H65+H107/CAF®01. Six weeks post final immunization, splenocytes of immunized mice were restimulated ex vivo with H65, H107, or single peptide epitopes $TB10.4_{71-88}$ or $ESAT\text{-}.6_{1-15}$ and total CD4 T cells expressing cytokines (TNF,IFN-γ,IL-2 and/or IL-17A) were enumerated by ICS. **(C)** The percentage of $I\text{-}A^d$:$TB10.4_{73-88}$ (red) or $I\text{-}A^b$:$ESAT\text{-}6_{4-17}$ (blue) tetramer binding $CD44^{high}$ CD4 T cells were identified in the lungs of non-vaccinated and BCG+H65+H107-vaccinated mice 27 (△) and 33 (○) days after aerosol Mtb infection. Symbols indicate individual animals. Box plots with whiskers indicate median, IQR, minimum and maximum values.

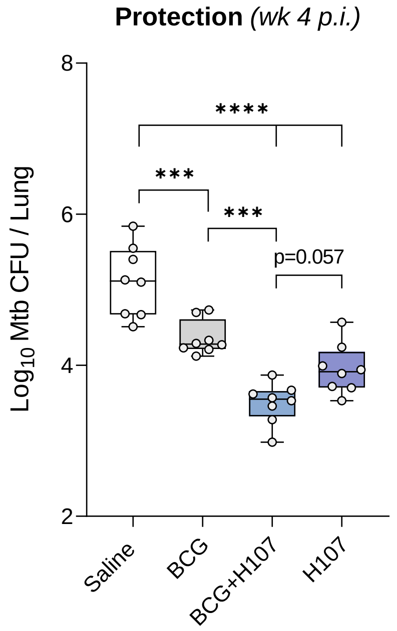


**Fig S5. Lung bacterial load in vaccinated and Mtb infected mice**

CB6F1 mice were immunized once s.c. with $0.5 \times 10^6$ CFU BCG Danish, three times s.c. with saline or 1 µg H107/CAF®01, or co-administered BCG+H107/CAF®01 as described previously (n=8). Six weeks later mice were infected with aerosol Mtb Erdman with a dose corresponding to 25-50 CFU per mouse. The bacterial burden was determined in the lungs four weeks post infection (p.i.). Boxplots indicate median, IQR, and minimum and maximum values. One-Way ANOVA with Tukey's multiple comparisons test. p-values; ***(p<0.001), ****(p<0.0001).

| | Strain | PPE68/ Rv3873 | EsxA/ Rv3675 | EspI/ Rv3876 | EspC/ Rv3615c | EspA/ Rv3615c | MPT64/ Rv1980c | MPT70/ Rv2875 | MPT83/ Rv2873 |
|---|---|---|---|---|---|---|---|---|---|
| | M.tuberculosis | +++ | +++ | +++ | +++ | +++ | +++ | +++ | +++ |
| | M.bovis | +++ | +++ | +++ | +++ | +++ | +++ | +++ | +++ |
| Early BCG strains (lacks RD1)# | BCG Russia | - | - | - | (+++) | (+++) | +++ | +++ | +++ |
| | BCG Japan | - | - | - | (+++) | (+++) | +++ | +++ | +++ |
| | BCG Moreau | - | - | - | (+++) | (+++) | +++ | +++ | +++ |
| | BCG Sweden | - | - | - | (+++) | (+++) | +++ | +++ | +++ |
| | BCG Birkhaug | - | - | - | (+++) | (+++) | +++ | +++ | +++ |
| Modern BCG strains (Lacks RD1 and RD2)$ | BCG Tice | - | - | - | (+++) | (+++) | - | (+) | (+) |
| | BCG Frappier | - | - | - | (+++) | (+++) | - | (+) | (+) |
| | BCG Pasteur | - | - | - | (+++) | (+++) | - | (+) | (+) |
| | BCG Danish | - | - | - | (+++) | (+++) | - | (+) | (+) |
| | BCG Glaxo | - | - | - | (+++) | (+++) | - | (+) | (+) |
| | BCG Prague | - | - | - | (+++) | (+++) | - | (+) | (+) |
| | BCG China | - | - | - | (+++) | (+++) | - | (+) | (+) |

### The RD1 region was lost during the attenuation of *M. bovis* (1908-21)
$ BCG substrains that were accuired after 1927 have in addition to RD1 lost the RD2 region and have a mutation in *sigK*

- gene not present on chromosome
+++ high protein expression
(+) low protein expression due to *sigK* mutation
(+++) high expression but the protein is not secreted due to a defect in the ESX-1 secretion system

**Table S1. Antigen expression of BCG substrains.**

Overview of the protein expression/secretion of the antigens comprising H107 (*71*).

| Name | Rv. No. | Protein length | Part included in H107 | Modifications |
|---|---|---|---|---|
| **PPE68** | Rv3873 | 368 | 155-219, GGS linker, 242-305 | Regions with homology to BCG removed **(72)** |
| **ESAT-6 / EsxA** | Rv3875 | 95 | 2-95 | No modifications |
| **EspI** | Rv3876 | 666 | 2-666 | K425Q (inactivation of ATP binding site) **(73)**<br>C537S, C590S (cysteine's replaced with serine's) |
| **EspC** | Rv3615c | 103 | 2-54 | C-terminal part removed for compatibility with ESAT-6 free IGRA **(33)** |
| **EspA** | Rv3616c | 392 | 2-131, 155-392 | Transmembrane / hydrophobic amino acids removed |
| **MPT64** | Rv1980c | 228 | 24-228 | Signal sequence omitted<br>C29S, C41S (cysteine's replaced with serine's) |
| **MPT70** | Rv2875 | 193 | 31-193 | Signal sequence omitted<br>C38S, C172S (cysteine's replaced with serine's) |
| **MPT83** | Rv2873 | 220 | 31-220 | Signal sequence omitted<br>C64S, C198S (cysteine's replaced with serine's) |

**Table S2. Overview of antigen modifications in H107.**
